# A linear ontogeny accounts for the development of naive, memory and tumour-infiltrating regulatory T cells in mice

**DOI:** 10.1101/2024.07.10.602914

**Authors:** Sanmoy Pathak, Thea Hogan, Sanket Rane, Yundi Huang, Charles Sinclair, Simon Barry, Larissa Carnevalli, Andrew Yates, Benedict Seddon

## Abstract

Foxp3^+^ Regulatory T cells (Treg) are a subset of CD4^+^ T cells that play critical functions in maintaining tolerance to self antigens and suppressing autoimmunity, regulating immune responses to pathogens and have a role in the pathophysiology of anti-tumoural immunity. Treg ontogeny is complex since they are generated following recognition of self antigens in the thymus during normal T cell development (thymic Treg), but are also induced from mature conventional T cells when activated by foreign antigen with appropriate additional cues (inducible Treg). How these distinct ontogenic pathways contribute to the maintenance and function of the mature Treg compartment in health and disease remains unclear. Here, we use a combination of fate mapping approaches in mice to map the ontogeny of Treg subsets throughout life and estimate rates of production, loss and self-renewal. We find that naive and effector/memory (EM) Treg subsets exhibit distinct dynamics but are both continuously replenished by de novo generation throughout life. Using an inducible Foxp3-dependent *Cre* fate reporter system, we show that naive Treg and not conventional T cells, are the predominant precursors of EM Treg in adults. Tonic development of new EM Treg is not influenced by foreign antigens from commensals, rather suggesting a role for self recognition. To investigate the ontogeny of Treg development in malignant disease, we used the same fate reporter systems to characterise the Treg infiltrate of three different model tumours. In all three cases, we found that Treg derived from pre-existing, EM Treg. Together, these results reveal a predominantly linear pathway of Treg development from thymic origin to EM Treg associated with pathophysiology of malignant disease, that is driven by self antigen recognition throughout.

## Introduction

Regulatory T cells (Treg) are a subset of CD4^+^ αβ T cells that have evolved to provide suppressive regulation of adaptive immune responses. Their suppressive function was first recognised in the context of autoimmune disease, where they play a vital role in maintaining the immune system in a state of active tolerance of self antigens (Saoudi et al., 1996b). It is now recognised that Treg also have a broader impact on normal immune function, normal adaptive immunity to pathogens (Belkaid and Tarbell, 2009), tissue homeostasis (Panduro et al., 2016) and in the development of malignant disease (Togashi et al., 2019). Adoptive T cell therapy has emerged as a potent means of specifically supplementing host immunity, and Treg are being considered in this context for treatment of autoimmune conditions (Amini et al., 2022; McGovern et al., 2017). In malignant disease, Treg are common constituents of tumour infiltrates, and in many instances are a biomarker of poor prognosis where their suppressive activity is thought to counteract beneficial tumour immunity (Peng et al., 2012; Saleh and Elkord, 2020; Schnellhardt et al., 2022). On this basis, therapies that target regulatory T cells or their function have been met with striking successes in the clinic, restoring anti-tumoural immunity (Pardoll, 2012; Ribas and Wolchok, 2018; Wolchok, 2021). However, such treatments are also often associated with toxicity due to loss of self tolerance and specifically targeting intra-tumoral Treg remains a major challenge. Knowledge of how Treg are generated and maintained throughout life is therefore critical for fully understanding their physiological functions both in the normal healthy immune system, and for designing and predicting the long-term impact of immunotherapies targeting regulatory T cell activity.

Treg are characterised by expression of the signature transcription factor Forkhead Box protein 3 (Foxp3) (Josefowicz et al., 2012) and recirculate through secondary lymphoid organs via the blood. They are a heterogenous population, both in terms of differentiation state and ontogeny, deriving from two principle sources. The first is during normal thymopoeisis of T cells. Treg develop in the thymus (Sakaguchi, 2004; Saoudi et al., 1996a) from precursors amongst CD4 single positive (SP) thymocytes and are termed thymic or natural Treg (tTreg hereon). The mechanisms by which these cells develop have been studied extensively and it is now clear that a co-stimulation dependent agonist selection event, mediated by recognition of self determinants, drives induction of Foxp3 in a CD4 SP T cell with appropriate specificity, with stable commitment critically supported by the cytokines IL-2 and IL-15 (Fu et al., 2012; Lio and Hsieh, 2008; Marshall et al., 2014; Ohkura et al., 2012; Tai et al., 2005; Tai et al., 2013). The second source of circulating Treg is through the induction of Foxp3 expression in conventional CD4^+^ T cells, termed inducible Treg (iTreg) (Kanamori et al., 2016). This pathway is characterised in vitro through activation in the presence of TGFβ(Chen et al., 2003), but has also been observed in vivo at intestinal sites in response to antigen challenge(Curotto de Lafaille et al., 2008; Ohnmacht et al., 2015) and in malignant disease(Weiss et al., 2012). In the latter case, tumours are thought to be potent stimuli of iTreg formation. Treg isolated from MC38 tumours express Nrp1 at low levels (Weiss et al., 2012), which has been used as a marker of iTreg. The extent to which iTreg contribute to the circulating Treg pool found in secondary lymphoid organs remains unclear, however.

In addition to these distinct ontogenic pathways, peripheral Treg exhibit considerable functional and phenotypic heterogeneity. They are composed of naive-like CD44^lo^CD62L^hi^ CCR7^hi^ cells and more differentiated subsets that display an activated CD44^hi^CD62L^lo^ effector/memory (EM) surface phenotype, additionally expressing Ox40, GITR, CD69, and exhibiting altered chemokine receptor expression (Campbell and Koch, 2011; Feuerer et al., 2009; Fisson et al., 2003; Huehn et al., 2004; Stephens et al., 2007). These states have been described during elicitation of iTreg following mucosal antigen challenge, and are associated with the generation and migration of Treg to mucosal sites (Akagbosu et al., 2022; Kedmi et al., 2022; Xu et al., 2018). Treg have been described that also express signature transcription factors associated with type 1 (Tbet)(Koch et al., 2009), type 2 (Gata3)(Yu et al., 2015), type 17 (RORgt)(Ohnmacht et al., 2015; Weiss et al., 2012) or T follicular helper responses (Bcl6)(Chung et al., 2011) and are specifically associated with regulation of the associated T helper responses.

While the developmental pathways that give rise to Treg have been studied extensively, the homeostatic mechanisms that are responsible for maintaining them are less clear. Basal Ki67 expression by Treg shows they undergo greater cell division than naive conventional T cells (Attridge and Walker, 2014), suggesting that self renewal contributes to Treg maintainance, as is true of conventional CD4^+^ memory T cells (van Leeuwen et al., 2009). Similar to naive conventional T cells, Treg require constitutive TCR signaling for their long-term survival and/or proliferation since genetic ablation of TCR expression results in loss of Treg, with a half life of approximately 5 weeks (Vahl et al., 2014). The role of thymic Treg generation in maintaining peripheral pools is also uncertain. tTreg appear to be present in the thymus throughout life, implying a role for replenishment of peripheral compartments with new cells. However, studies of Rag2-EGFP mice reveal that as mice age, an increasingly large fraction of these cells are mature Treg that have recirculated back to the thymus (Cowan et al., 2016; Thiault et al., 2015), and it has been suggested that recirculated Treg in the thymus compete for IL-2 and directly inhibit intrathymic Treg development(Thiault et al., 2015). We therefore know little about the extent to which thymic Treg generation contributes to maintenance of peripheral pools throughout life, and how EM Treg are maintained if thymic output is indeed shut down. In this context, it is also unclear how the origin and specificity of EM Treg changes over the life course, and whether they derive from naive Treg precursors, or are instead iTreg derived from conventional precursors.

In total, considerable uncertainty remains around how peripheral Treg subsets are generated and maintained, the relative role of thymic Treg generation to their replenishment throughout life, and how their homeostatic dynamics contribute to the life-long maintenance of this critical population. In the present study, we employed a variety of fate mapping systems to dissect the ontogeny and temporal dynamics of Treg generation and persistence across the mouse lifespan, and in the setting of malignant disease.

## Results

### Thymic Treg comprise both *de novo* generated and recirculated mature cells

To understand the cellular mechanisms governing the maintenance of the regulatory T cell compartment over the lifespan of a mouse, we first generated and analysed two independent datasets. The first was an ontogenic analysis of T cell compartment size and phenotype in normal mice throughout their life course. The second dataset came from experiments that employed a busulfan drug-conditioned bone marrow chimeric mouse model that permits the contribution of de novo haematopoesis to mature blood cell compartments to be measured and quantified over time; so-called temporal fate mapping (Hogan et al., 2015). In this model, conditioning hosts with busulfan specifically ablates the haematopoetic stem cell (HSC) compartment, allowing its reconstitution with congenically labelled stem cell progenitors. Importantly, mature haematopoetic lineages are unaffected by this conditioning, therefore allowing the infusion of new donor origin cells into existing replete compartments to be detected and measured over time. The kinetics and extent of replacement of existing cells with new cells in any given cellular compartment are rich in information about the dynamics underlying its maintenance (Gossel et al., 2017; Hogan et al., 2015; Hogan et al., 2019; Rane et al., 2022). Cohorts of chimeras were generated by treating B6.CD45.1 host mice with drug and the HSC compartment reconstituted 24h later with donor B6.CD45.2 bone marrow. In order to assess potential host age effects upon Treg homeostasis, chimeras were generated with hosts of a range of ages (6-8wks, 8-10wks, 10-12wks and 12-25wks). We then analysed Treg phenotype in thymus, lymph nodes and spleen of chimeras between 14 and 300 days post-BMT. The extent of HSC reconstitution (donor chimerism) was assessed by measuring the donor fraction amongst CD4 CD8 double positive (DP) thymocytes, which stabilizes within 5 weeks after BMT and serves as an accurate proxy of average HSC chimerism across all bone marrow sites (Verheijen et al., 2020). Following busulfan treatment, donor fractions amongst DP thymocytes were between 70-100% (Fig. 1A). Normalising the observed donor fraction in any downstream population to the equilibrated donor fraction in early DP-stage thymocytes accurately reflects the extent to which that population has been replaced through de novo haematopoesis since BMT.

**Figure 1.**
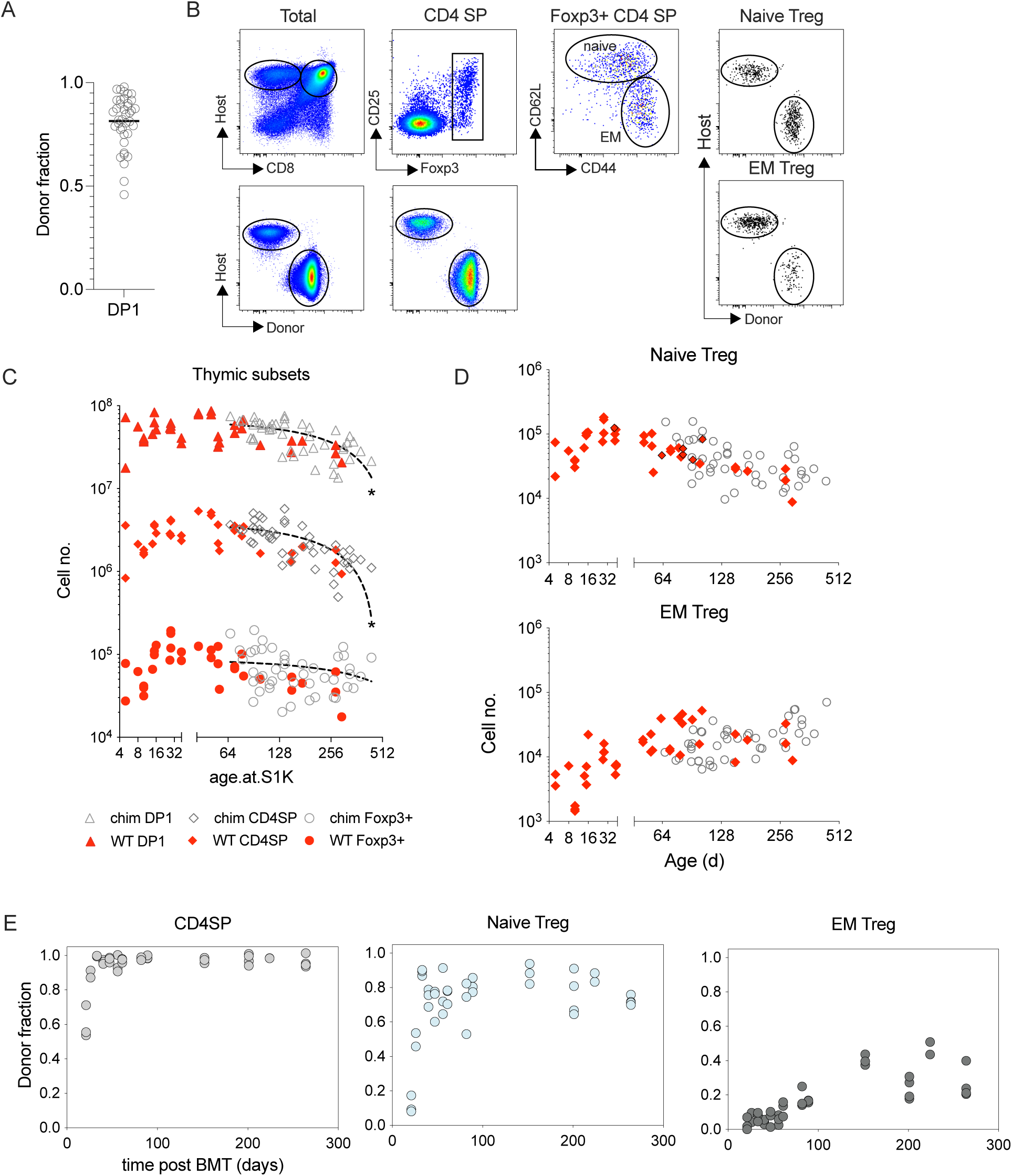
Thymic development of regulatory T cells. Busulfan chimeras were generated using B6.CD45.1 hosts and B6.CD45.2 donors (See Methods) and mice analysed at various times after BMT. (A) Scatter plot of donor chimerism in DP1 thymocytes of all chimeras analysed. (B) Gating strategy to identify naive (CD62L^hi^CD44^lo^) and EM (CD62L^lo^CD44^hi^) Foxp3^+^ Treg in thymus, and donor (CD45.2) vs host (CD45.1) composition therein. (C) Scatter plots shows total number of DP1, CD4 single positive (CD4SP) and Treg (CD4SP Foxp3^+^) subsets isolated from thymus of chimeras (black open) and untreated WT control mice (red diamonds), with mouse age. (D) Scatter plots of total numbers of naive and EM Treg from chimeras and WT control, by host age. (E) Scatter plots of normalised donor fraction at different times postBMT amongst CD4SP, naive Treg and EM Treg subsets. Data are pooled from multiple independent batches of chimeras (n=45) and WT controls (n=34). For linear regression lines, * p < 0.0001 that slope is non-zero.

We first analysed the dynamics of Treg development in the thymi of both WT and busulfan chimeras (Fig. 1B). In mice, the thymus actively produces new T cells throughout life. However, both thymic size and output peaks at around 6 weeks of age, after which the thymus undergoes a slow but steady decline in both size and productivity, termed thymic atrophy (Boehm and Swann, 2013). Measuring total numbers of DP and CD4 single positive (CD4SP) thymocytes in control mice and chimeras confirmed the expected peak and subsequent decline in numbers (Fig. 1C). We have previously estimated that these populations wane exponentially with age, halving in size approximately every 150d (Hogan et al., 2015). In contrast, total Foxp3^+^ Treg numbers in the thymus showed no significant decline with age (Fig. 1C).

Previous studies using Rag2-EGFP reporter mice show that a substantial fraction of Treg in the thymus are mature GFP-negative cells that have recirculated from the periphery (Thiault et al., 2015) in a CCR7-dependent manner (Cowan et al., 2016). We confirmed the presence of thymic Treg with both naive and EM phenotypes (Fig. 1B). However, the composition of this heterogeneous population shifted over time; naive Treg numbers declined significantly with age (Fig. 1D), while EM Treg numbers increased, reaching a plateau around 9 weeks of age, but remained relatively stable thereafter (Fig. 1D). Importantly, we saw no difference in the numbers or naive/memory composition of thymic Treg between WT and busulfan chimeric mice. To confirm the ontogeny and source of these intrathymic populations, we next analysed the infusion of donor-origin cells into these subsets over time, in busulfan chimeric mice. The normalised donor fraction within CD4SP thymocytes rapidly reached 1 within a few weeks post BMT, indicating that this population had turned over completely (Fig. 1E). Naive thymic Treg also underwent a rapid replacement by donor progeny, but failed to achieve complete replacement, reaching a plateau of ∼0.8 (Fig. 1E). In contrast, donor-origin EM Treg were not readily detectable until almost 100 days post BMT, even though numbers of memory Treg in the thymus peaked by ∼d63 of age. The presence of host-derived naive and memory Treg within the thymus validates the conclusion that these Treg are not de novo generated but recirculated from the peripheral mature Treg compartments. Previous studies have suggested that mature Treg that recirculate in the thymus compete for IL-2 and thereby impair de novo Treg generation that is IL-2 dependent (Thiault et al., 2015). To quantify any inhibition in our experiments, we needed knowledge of the donor:host compositions of both recirculating and newly-generated naive T cells. We therefore analysed the dynamics of Treg turnover in the periphery.

### Circulating Treg that migrate between lymphoid organs are heterogeneous in composition but in equilibrium between lymphoid sites

We analysed the dynamics of peripheral naive and EM Treg subsets in lymph nodes and spleen of both WT mice and busulfan chimeras, measuring donor/host chimerism and Ki67 expression as an indicator of cell division (Fig. 2A). From birth, total numbers of naive Treg rose rapidly in young mice, reaching maximal levels as early as 5 weeks of age. Numbers were maintained for several months after which a gradual decline was evident (Fig. 2B). In contrast, memory Treg were scarce in young mice, but gradually increased in number until 9-10 weeks of age, and continued to accumulate more slowly thereafter (Fig. 2C). Correspondingly, naive:EM Treg ratios in both LN and spleen were high early in life but declined progressively, with memory Treg exceeding naive Treg numbers at around 8 months of age (Fig. 2D). Analysing the kinetics of donor cell infusion in busulfan chimeras also exposed distinct dynamics. A steep rise in donor cell infusion was observed amongst naive Treg, but the population as a whole failed to reach complete replacement, with the normalized donor fraction reaching a plateau by around d100. In contrast, donor infusion into the EM Treg pool was much slower and reached a far lower level of replacement.

**Figure 2.**
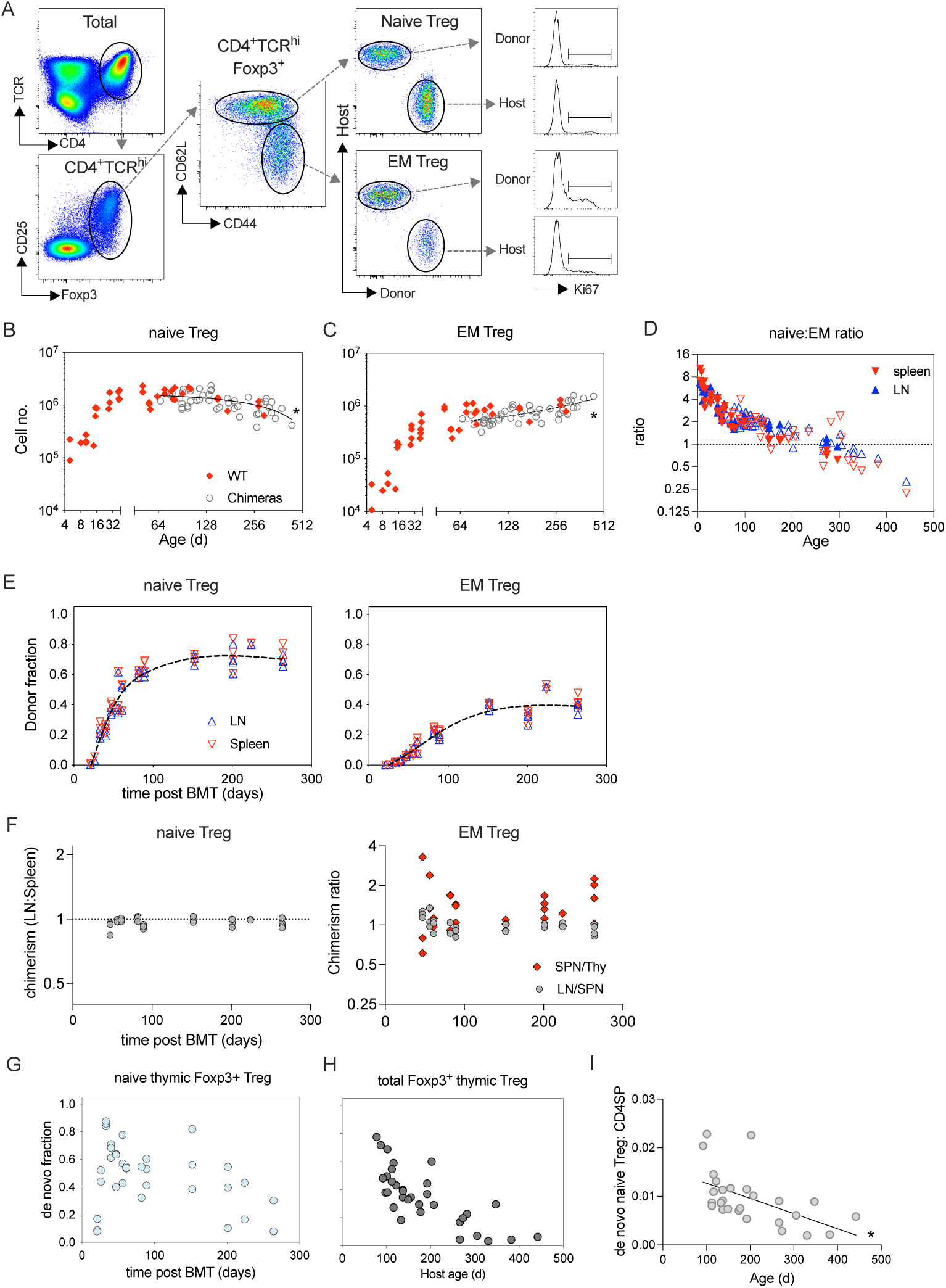
Mapping the developmental dynamics of regulatory T cell subsets in busulfan chimeras. Cells from lymph node and spleen from busulfan chimeras and WT mice described in figure 1 were analysed. (A) Gating strategy to identify naive (CD62L^hi^CD44^lo^) and EM (CD62L^lo^CD44^hi^) Foxp3^+^ Treg in peripheral lymphoid organs, their donor (CD45.2) vs host (CD45.1) composition therein, and gates used to measure Ki67 expression. (B-C) Scatter plots are of total numbers of naive (B) and EM (C) Treg recovered from lymph node and spleen of chimeras and control WT mice of different ages. Lines show simple linear regression fits to data. (D) Scatter plot of ratio of naive to EM Treg in lymph node and spleen of WT (filled symbols) and busulfan chimeras (empty symbols) at different host ages. (E) Scatter plots of normalised donor fraction in lymph node and spleen of busulfan chimeras at different times post BMT for naive and EM Treg. (F) Scatter plots showing of ratios of donor chimerism in naive Treg between Lymph node : Spleen, and in EM Treg between Spleen:thymus and LN:spleen. Panels G-H; The estimated fractions of de novo developed Treg amongst (G) total naive thymic foxp3^+^ cells and (H) total (naive and memory) Foxp3^+^ thymic Treg. (I) De novo generated naive Treg as a fraction of total CD4SP thymocytes. For linear regression lines, * p < 0.0001 that slope is non-zero.

Our enumeration of Treg subsets assumed that populations in lymph node and spleen represented intimately mixed, freely recirculating populations of cells in equilibrium. If true, the chimerism in lymph node and splenic populations at any given time should be closely matched, regardless of the dynamic changes observed over time, following BMT. Calculating ratios of donor chimerism between LN and spleen of individual mice confirmed this view. Ratios amongst naive and memory Treg both remained around 1 throughout the time course of Treg renewal, independent of the extent of donor chimerism (Fig. 2F). The chimerism of EM Treg within the thymus was also very similar to that in spleen, indicating that these cells recirculate and are in dynamic equilibrium with populations in secondary lymphoid organs.

### *De novo* production of naive thymic Treg continues with age

The failure of donor naive Treg to completely replace host naive Treg populations in the thymus strongly suggested that some fraction of naive thymic Treg had also recirculated from the periphery, in addition to the recirculation of EM Treg. To estimate this fraction, we assumed that donor/host composition of recirculated naive Treg in the thymus would match that of peripheral naive Treg, in any given individual. The additional fraction of donor naive Treg observed in the thymus would therefore represent de novo generated naive thymic Treg. We estimated this additional fraction (Fig. 2G) and showed that at early timepoints after BMT, most naive Treg are de novo generated, but over time their representation reduces. Earlier studies analysed how the fraction of total Foxp3^+^ thymic Treg that expressed Rag2-GFP, changed with age, revealing a dramatic collapse in GFP^+^ fraction with age. Estimating this same fraction from our data, assuming all memory Treg were recirculated cells, resulted in very comparable results to this earlier study (Fig 2H). However, the increasing numbers of Treg that recirculate to the thymus with age, together with the natural atrophy of the thymus with age, obscures the extent to which de novo Treg generation changes with age. To explicitly measure this change, we calculated the ratio of estimated de novo naive Treg to the size of the total CD4 SP precursor pool. In young mice, de novo generated naive Treg are ∼0.01 of the total CD4 SP compartment. However, this fraction undergoes a modest but significant reduction with age (Fig. 2I). Nevertheless, this analysis reveals that thymic production of naive Treg does continue even in aged mice and in the presence of ever-increasing numbers of mature Treg that have recirculated back to the thymus.

### Thymic export and self renewal both contribute substantially to the homeostasis of naive Treg

This empirical analysis revealed that the naive and EM Treg compartments exhibited distinct shifts in their size and rates of infusion of new cells with age. To better understand and quantify the cellular processes responsible for these dynamics, we used mathematical modelling to test different models of the influx, self-renewal and loss that underlie them. We first analysed the naive Treg compartment, comparing the support for two simple models (Fig. 3A); a ‘homogeneous’ model in which all naive Treg exhibit identical dynamics, and an ‘incumbent’ model, in which host naive Treg are heterogeneous, comprising a subset that shares the same homeostatic dynamics as newly generated donor thymic Treg, and a subset that is resistant to displacement by new donor Treg and may exhibit distinct rates of division and loss. Previously we found support for this ‘first-in, last-out’ structure within conventional naive T cells (Hogan et al., 2015; Rane et al., 2022; Rane et al., 2018). An important feature of the data that the models needed to explain was the failure of donor cells to achieve complete replacement of host cells. The homogeneous model predicts complete turnover of a population given sufficient time and consistent influx of new cells, which we have observed in other lymphocyte populations in busulfan chimeric mice (Verheijen et al., 2020). Therefore, it can only explain incomplete turnover if the steady decline in de novo generation of naive Treg, which we and others (Thiault et al., 2015) have demonstrated, is sufficiently fast. Incomplete replacement is also explained naturally by the presence of incumbent, host-derived naive Treg, established before BMT and which persist independently of newly generated cells.

**Figure 3.**
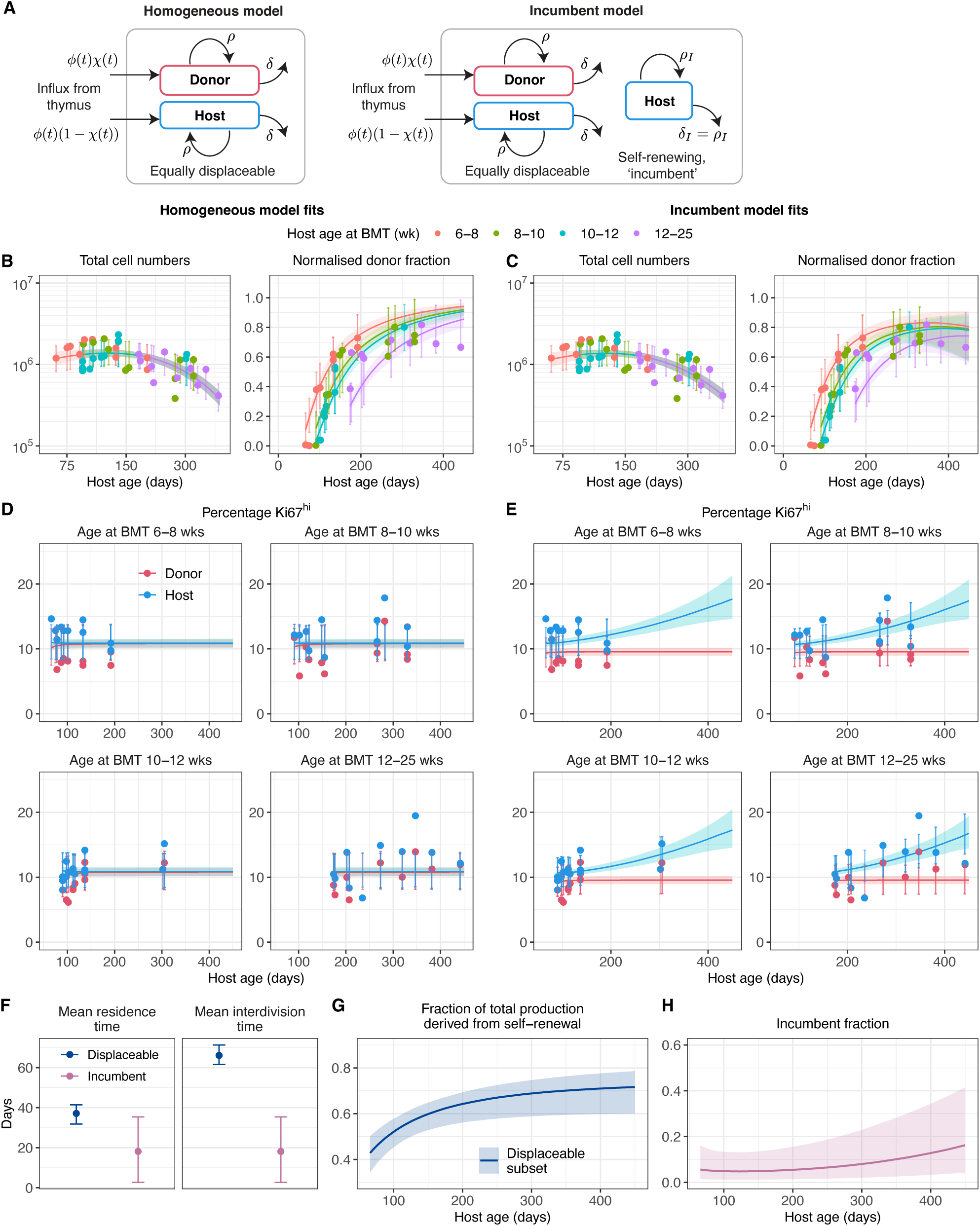
Naive Treg are heterogeneous in their homeostatic dynamics. (A) Schematics of the homogeneous and incumbent models of naive Treg homeostasis. (B,C) Best fits of the homogeneous and incumbent models, respectively, to the numbers and normalised donor fraction of naive Treg cells over time. (D,E) The two models’ fitted trajectories of Ki67 expression within donor and host cells. Mice were grouped by age at BMT, in weeks; 6-8 (n=10), 8-10 (n=11), 10-12 (n=9) and 12-25 (n=9). (F) Mean residence and interdivision times of naive Treg in the incumbent model, with 95% confidence intervals. (G) The estimated proportion of daily production of naive Treg that derived from self-renewal rather than de novo production. (H) The estimated proportion of Treg that were incumbent (host-derived, established before BMT). Shaded regions indicate 95% confidence envelopes.

We fitted each model simultaneously to the time courses of total numbers of naive Tregs, their host/donor composition, and the levels of Ki67 expression among donor and host cells (Fig. 3B). We assumed that the influxes of host and donor cells into the naive Treg compartment were proportional to the numbers of their respective de novo generated thymic naive precursors, the time courses of which we described empirically (Fig. S1). While both models captured the changes in the total numbers of naive Treg with age (Figs. 3B and 3C, left panels), the homogeneous model could not account for the incomplete donor infusion (Fig. 3B, right panel), even given the waning production of new naive Treg by the thymus (Fig. 2I, S1). This model also failed to account for the higher frequency of Ki67 expression among host cells than among donor cells (Fig. 3D). In contrast, the incumbent model provided better descriptions of these key observations (Fig. 3E), indicating that the naive Treg compartment is heterogenous with respect to its propensity for replacement. This simple model of heterogeneity predicted that displaceable naive Treg had a mean residence time of around 40 days and divided on average every 60 days (Fig. 3F), whereas incumbent Treg, which were assumed to be stable in numbers, divided and were lost every 20 days. The displaceable naive Treg population was sustained by influx from thymic precursors throughout life. However, the contribution of self renewal becomes increasingly important as mice aged, a contribution estimated to increase to close to 70% in mice over a year old as precursor numbers declined. In tandem, incumbent cells made up an increasing proportion of the population with age (Fig. 3H). This kinetic explains the diverging levels of Ki67 expression among donor and host cells with age (Fig. 3D).

### EM Treg derive predominantly from naive Treg rather than conventional T cell subsets

Next, we extended our modeling approach to investigate the ontogeny and homeostasis of memory Treg. While our data and others clearly suggest that naive peripheral Treg derive directly from thymic naive Treg, the extents to which memory Treg derive from naive Treg or from conventional T cell subsets is unclear. We therefore explored models in which the precursors of memory Treg are either naive peripheral Treg, or are instead iTreg, deriving from a Foxp3-negative conventional T cell subset. We gave consideration to each of naive (CD44^lo^CD62L^hi^), central memory (CD44^hi^CD62L^hi^) and effector memory (CD44^hi^CD62L^lo^) subsets as potential precursors amongst conventional CD4^+^ T cells. Irrespective of the precursor, we reasoned that differentiation into memory Treg occurs through activation and cell division. Therefore we explored a simple two-compartment model in which precursors enter the memory Treg compartment via a ‘fast’ subset that turns over quickly before differentiating to a more quiescent ‘slow’ state with lower rates of division and death (Fig. 4A). Previously we found support for similar behaviour within circulating conventional memory CD4 T cells (Gossel et al., 2017; Hogan et al., 2019). To define the influx into host and donor memory Treg in this model, we used empirical descriptions of the time courses of the numbers and donor chimerism of candidate precursors of memory Treg (Fig. S2). By fitting this model to the total numbers of memory Treg as well their donor/host composition and Ki67 levels, we found support for naive Treg as their primary precursor. However, the statistical support for this pathway over the others was modest with a LOO-IC weight of 52%, compared with 26%, 13.5% and 8.5% for TEM, TCM and naive Tconv subsets respectively. Nevertheless, this best fitting model was able to explain the changes of memory Treg numbers with age (Fig. 4B) and the kinetics of their replacement by donor cells (Fig. 4C). The model also explained the increased expression of Ki67 amongst donor cells relative to host cells at early times after BMT (Fig. 4D), when donor cells are more prevalent within the more recently recruited ‘fast’ subset. Self-renewal did not compensate for the loss of fast cells (Fig. 4E), indicating a role for influx in driving the accumulation of EM Treg during the first year of life; this comprised approximately 1% of memory Treg are per day, although this proportion declines with age (Fig. 4F). We estimated that cells persist in the ‘fast’ state for about a week before transitioning to slow cells, consistent with the normal dynamics of T cell activation, and that ‘slow’ EM Treg divide and die on average roughly every 40 days (Fig. 4E). From this similarity in rates of self-renewal and loss we predict that cohorts of slow EM Treg can persist dynamically (that is, at a population level) for many months. The model also explains the increase in total numbers of EM Treg during the first year of life (Fig. 4B); during this period, even though the total rate of influx of new memory from its precursor is declining, it still outstrips the net rate of loss of fast cells, continually boosting their numbers. This accumulation in turn drives the accumulation of slow memory cells, explaining the decline in Ki67 among EM Treg in bulk across the same period (Fig. 4D). See Supporting Information S5 for details.

**Figure 4.**
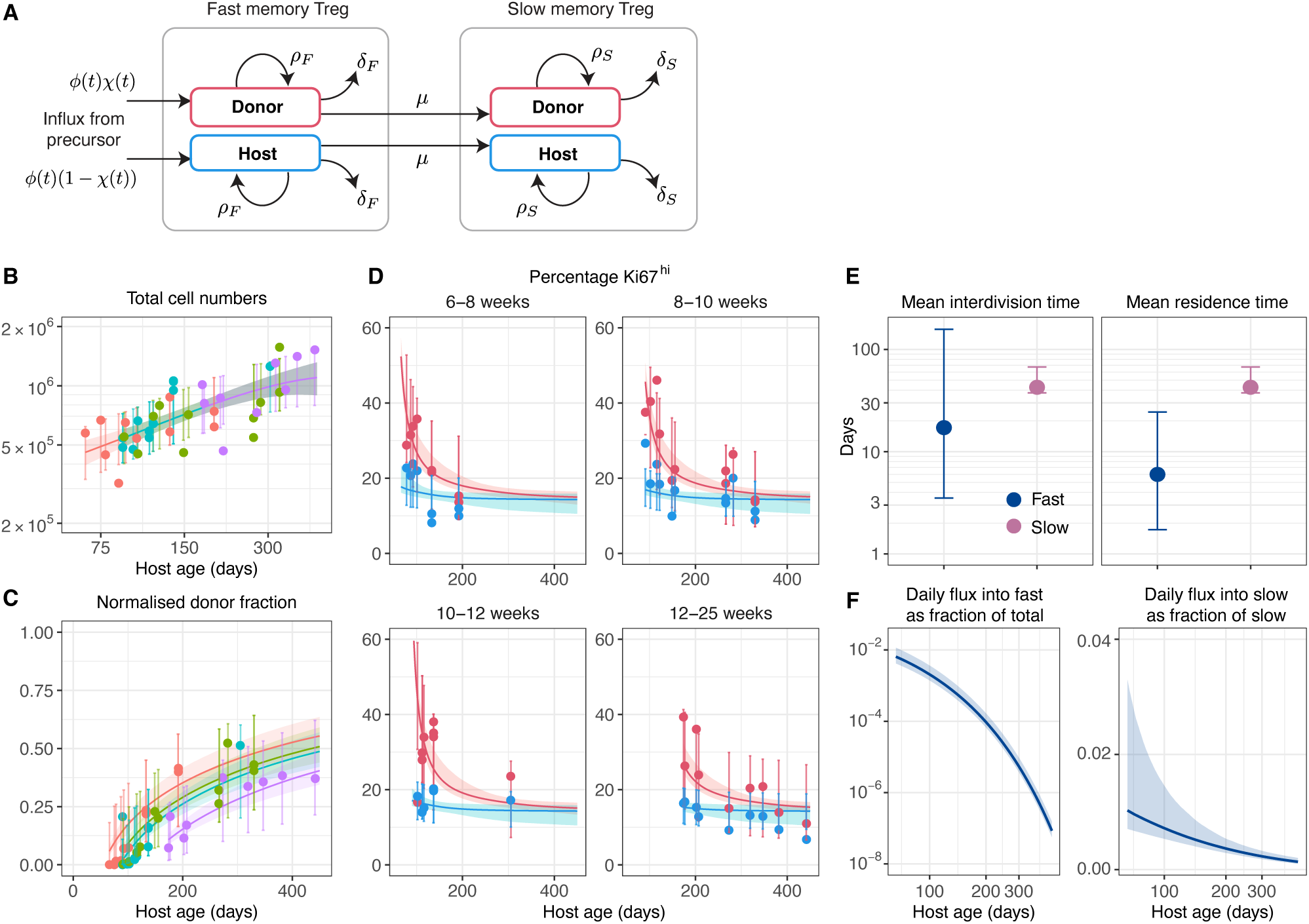
Memory Treg are constitutively replenished from naive Treg and transition from fast to slow turnover following their de novo generation. (A) Schematic of the two-compartment model of memory Treg homeostasis. (B-D) The fitted time courses of the total numbers of memory Treg (B), donor fraction (C), and levels of Ki67 (D) in busulfan chimeric mice, grouped by age at BMT (6-8wks, n=10; 8-10wks, n=11; 10-12wks, n=9; and 12-25wks, n=9). (E) Estimated mean interdivision times and residence times (mean time before death or onward differentiation) of fast and slow memory Treg. (F) Daily proportional influx of fast and slow subsets by newly generated memory.

### Foxp3-CreER temporal fate mapping supports naive Treg as the primary source of EM Treg

To test the conclusion that naive Treg are the precursors of recirculating memory Treg, we used a *Foxp3^GFP-CreERT^ Rosa26^RmTom^* fate reporter strain (Foxp3FR hereon). In this mouse, a tamoxifen (TAM)-inducible GFP-CreERT construct is expressed downstream of the endogenous Foxp3 locus. Induction of Cre activity by TAM administration results in the induction of permanent and heritable mTom reporter expression by cells currently expressing Foxp3. A single administration of TAM induced mTom in approximately 75% of Foxp3^+^ cells (Fig. 5A). Further TAM treatments did not substantially increase this frequency and so we continued with the minimal intervention of a single treatment. We followed the phenotype of Treg in Foxp3FR mice for 130 days after TAM administration, during which the fraction of Foxp3^+^ CD4^+^ T cells labelled with mTom declined gradually (Fig 5A). Analysing Foxp3-GFP and CD25 expression by mTom-labelled CD4^+^ T cells confirmed that mTom induction was strictly specific to the Foxp3^+^ Treg lineage. Our analysis also demonstrated that the Treg lineage fate was stable over time, since labelled cells retained a CD25^hi^Foxp3^+^ phenotype throughout the chase period (Fig. 5A), consistent with earlier studies (Rubtsov et al., 2010).

**Figure 5.**
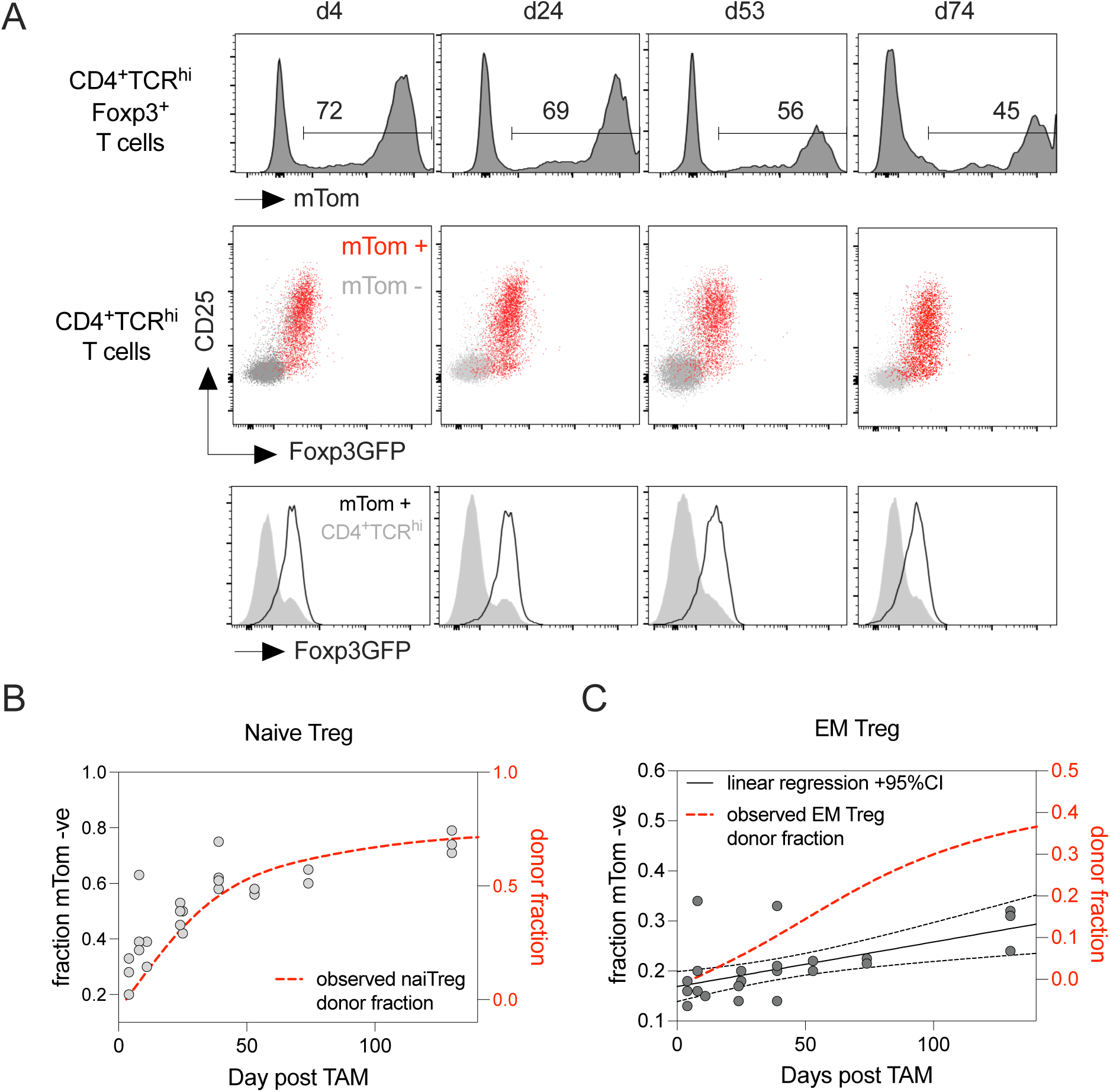
Foxp3-FR fate mapping implicates naive Treg as the source of de novo EM Treg. Foxp3-FR mice were treated with a single feed of 2mg of tamoxifen and cohorts of mice taken at different times after treatment. (A) Histograms (top row) show mTom reporter expression at the indicated days after treatment by CD4^+^TCR^hi^ Foxp3^+^ cells from lymph nodes. Scatter plots show CD25 vs Foxp3EGFP expression by mTom+ve and mTom-ve CD4^+^ T cells from lymph nodes. Histograms (bottom row) are of Foxp3-EGFP expression by mTom^+^ CD4^+^TCR^hi^ T cells or total CD4^+^TCR^hi^ cells from lymph nodes at different times after treatment. (B-C) Scatter plots are of fraction of naive (B) or EM (C) Foxp3^+^ Treg that are mTom -ve, calculated from total numbers of Treg recovered from LN and spleen. Splines (long dashes) are best fit lines of normalised donor fractions observed in busulfan chimeras in either naive or EM Treg as shown in figure 2E. A linear regression line with 95% confidence intervals is applied to mTom-ve fraction of memory Treg. Slope deviation from 0, p = 0.013.

The decline in the frequency of mTom+ cells among naive Treg after treatment was expected, as our analysis of busulfan chimeras showed that naive Treg are replaced constitutively by newly generated cells from the thymus (Fig. 2C). The kinetic of dilution of mTom+ naive Treg following TAM treatment of Foxp3FR mice might then be expected to mirror that of the replacement of host by donor cells in the busulfan chimeras. To explore this, we noted that following TAM treatment, approximately 20% of naive Treg were mTom-negative (Fig. 5B). We fitted a smooth curve (spline) to the kinetic of host cell replacement in busulfan chimeras (Fig. 2E), and overlaid this curve on the time course of the observed mTom-negative fraction, aligning them at the 20% level (Fig. 5B, dashed line). The subsequent kinetics were in good agreement, validating the conclusions we drew from the busulfan chimeric mice.

To test our assertion that naive Treg, rather than naive conventional T cells, are the primary precursors of EM Treg, we followed mTom expression among memory Treg following TAM treatment of Foxp3FR mice. Because Foxp3^+^ Treg but not conventional T cells were labelled with mTom (Fig. 5A), we reasoned that the kinetics of dilution of mTom-expressing EM Treg would reveal the extent to which new cells derived from a labelled (i.e. pre-existing naive Treg) or unlabelled (i.e. conventional T cell) precursor. Specifically, if precursors are unlabelled, the loss of mTom-expressing memory Treg is expected to be an immediate and acute event. In contrast, if the precursors were mTom labelled, new EM Treg would also be labelled, and thus the kinetic of loss of mTom-expressing EM Treg would be determined instead by the loss of labelled cells among naive Treg precursors. In busulfan chimeras we saw that almost ∼40% of EM Treg were replaced by new influx between d25 and d150 post BMT. However, in TAM-treated Foxp3FR mice, we did not observe any significant decline in the mTom+ fraction within EM Treg over a similar timeframe (Fig. 5C), indicating that cells entering the memory Treg pool in this interval were also rich in mTom+ cells. The most parsimonious explanation for these dynamics is that naive Treg are indeed the predominant or exclusive source of EM Treg in the steady state.

### EM Treg development is driven by self recognition independently of microbiome

Our data from busulfan chimeras demonstrated that memory Treg are continuously generated throughout life, via a Ki67^hi^ intermediate, strongly suggestive of an activation linked cell division process. This led us to consider the nature of the antigens driving EM Treg generation. Given that naive Treg are selected on self antigens in the thymus, it would be logical that recognition of self antigens in the periphery could result in transition to an activated phenotype. Alternatively, interactions with foreign antigens from the microbiome could also drive EM Treg generation, as has been described in the gut mucosa (Akagbosu et al., 2022; Kedmi et al., 2022; Xu et al., 2018). To explore these possibilities, we took advantage of a natural experiment that occurred when our mouse colony was relocated from a facility using open cages and conventional diets and tap water (which we refer to for brevity as “dirty”) to a new “clean” facility in which mice were housed in individually ventilated cages and fed sterile irradiated diet and water. The data presented in Figs.1-2 were generated from WT mice and busulfan chimeras held in the clean facility. However, similar experiments were conducted in the dirty facility prior to relocation. Our previous analyses of these data showed that the memory compartments of conventional T cells are substantially enlarged in mice held in dirty conditions, and that the natural microbiota play a critical role in establishing memory compartments in mice (Hogan et al., 2019). As such, the ratio of memory:naive populations in dirty hosts was elevated throughout life, compared with clean hosts (Fig. 6A). We therefore performed a meta-analysis to examine the steady state generation of naive and EM Treg in busulfan chimeras housed in dirty and clean settings. Comparing the normalised donor fractions in the two sets of chimeras revealed similar trajectories of donor cell infusion into both naive and EM Treg subsets (Fig. 6B). Similarly, the kinetics of total donor Treg numbers that appeared following BMT confirmed this view, with absolute numbers of naive or EM donor Treg reaching identical levels in both clean and dirty hosts (Fig. 6C). Together these results demonstrate that the tonic generation of EM Treg occurred independently of host microbiota in our systems.

**Figure 6.**
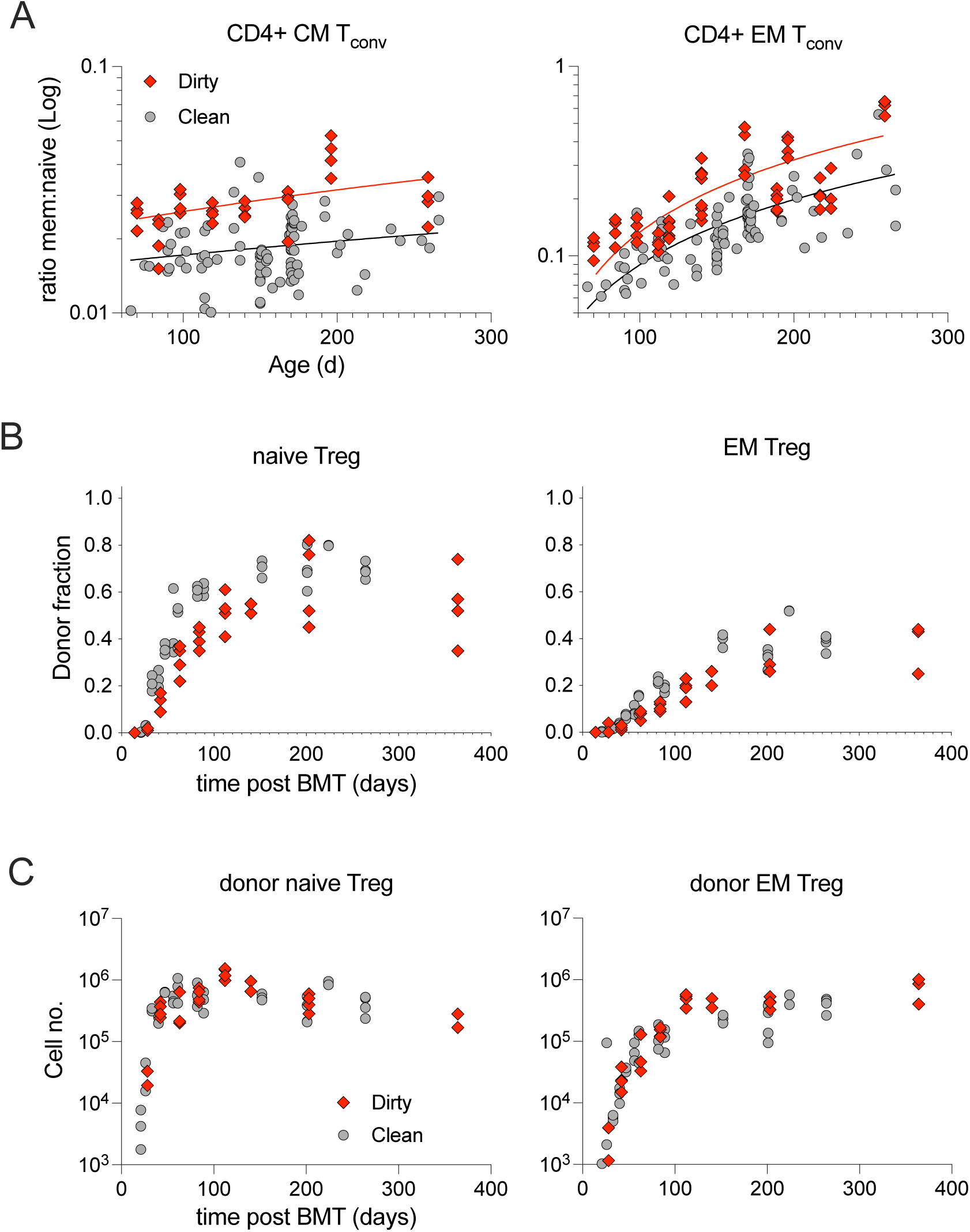
Tonic memory Treg generation is insensitive to environmental antigens. Data from busulfan chimeras detailed in fig. 1 (grey symbols, clean), generated and analysed at the UCL Comparative Biology Unit, were compared with chimeras generated in an identical manner three years previously, at the MRC National Institute for Medical Research (red symbols, dirty). In this meta analysis, Treg were identified by CD4^+^CD25^+^ gate throughout, since earlier study did not include Foxp3 intracellular stain. (A) Ratio of memory:naive CD4^+^ T_conv_ cells calculated from total numbers of subpopulations recovered from lymph nodes and spleen from different experiments, for central (CM) and effector memory (EM) cells. Lines are linear regressions to data. (B) Comparison of normalised donor fractions (B) and absolute cell numbers of donors cells (C) for naive and EM Treg.

### Tumour infiltrating Treg derive from existing circulating EM Treg

Our results suggest that self-recognition drives a linear pathway of development of naive and EM Treg throughout life. We next asked how these dynamics applied in the context of malignant disease setting. Previous studies of MC38 tumours in mice characterise infiltrating Treg as being induced from conventional T cells and are identified by their low expression of the marker Nrp1 (CD304)(Weiss et al., 2012). We therefore examined the lymphocytic infiltrate of MC38 tumours engrafted into our different fate reporter models to more directly characterise the ontogeny of tumour infiltrating Treg.

We first used the Foxp3FR reporter strain to determine whether Treg in TME derived from pre-existing Foxp3^+^ Treg, or were induced from conventional T cells, following tumour grafting. Foxp3FR mice were treated with TAM in order to pre-label existing Treg, and then engrafted with MC38 cells. Tumours were allowed to grow to a maximum permitted size of 15mm, typically achieved by day 17, after which hosts were culled. Analysing cellular composition of tumour masses revealed the anticipated lymphocytic infiltrate of T cells (Fig. 7A). CD4^+^ T cells in the TME included a substantial fraction of Foxp3^+^ Treg that exhibited an activated EM phenotype with elevated expression of PD1 and Ki67 relative to naive and EM Treg from LN or spleen (Fig. 7B). Consistent with previous reports (Weiss et al., 2012), Nrp1 expression levels were modestly reduced on tumour infiltrating Treg (Fig. S3). Examining Foxp3 reporter expression by Treg in TME revealed that in all cases a large proportion of Treg recovered from tumours expressed mTom, indicating that they arose from Treg labelled prior to tumour engraftment. Furthermore, the frequency of mTom expression by Treg in tumour was similar to that of Treg from lymphoid tissues of the same host (Fig. 7B). Over the course of the experiment, mTom frequencies in naive and EM Treg compartments of LN and spleen diverged, as mTom-negative cells started to replenish the naive Treg pool and percolated into memory. A closer comparison of mTom frequencies in the different compartments revealed that the mTom fraction among tumour-infiltrating Treg was higher than that among naive Treg, and more closely matched that of EM Treg in lymphoid organs, suggesting that tumour-infiltrating Treg derive from circulating EM Treg. To explore this ontogenic relationship further, we undertook temporal fate mapping of Treg in the TME to assess their origin. To do this, we exploited our earlier observations in busulfan chimeras, that donor naive and donor EM Treg accumulated with highly distinct kinetics (Fig. 2E). A concordance of the donor fraction of Treg in TME with either naive or EM Treg would suggest strongly that they share a common precursor. Chimeras were generated as described earlier and were engrafted with MC38 cells at 8 weeks post BMT. At this time, the donor fraction among naive Treg was ∼0.4 but was much lower (∼0.1) among EM Treg. Measuring the donor fraction among Treg from tumours revealed low chimerism, closely resembling that of circulating EM Treg in lymphoid organs (Fig. 7C). These data strongly suggest that Treg that infiltrate MC38 tumours derived from pre-existing memory phenotype Treg.

**Figure 7.**
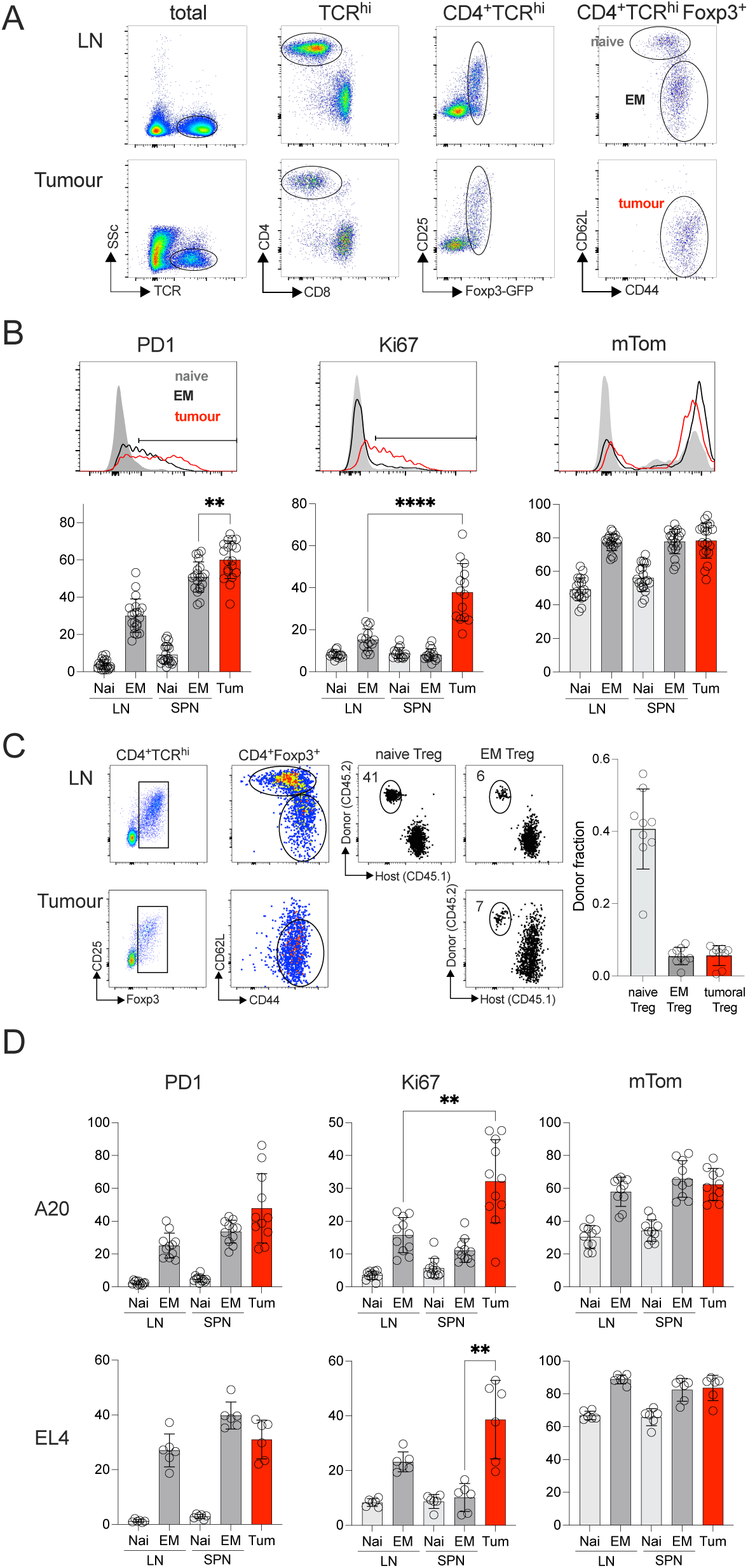
Intratumoral Treg derive almost exclusively from pre-existing circulating EM Treg. (A-B) Foxp3-FR mice were treated with a single feed of tamoxifen and 14 days later, engrafted with MC38 tumour cells. Mice were culled when tumour size reached maximal permitted size (15mm), between 15-20d post engraftment. (A) Density plots show TCR, CD4 and Foxp3 gates used to identify Treg in lymph node (LN) and tumour of host mice, and their naive vs EM composition. (B) Histograms and bar charts show representative and summarised expression of PD1, Ki67 and mTom by naive and EM Treg in LN and spleen, compared with tumour infiltrating Treg. (C) Busulfan chimeras were generated as described in figure 1. Hosts were engrafted with MC38 cells ∼8 weeks post BMT. Mice were culled when tumour size reached maximal permitted size, and phenotype of Treg in lymph nodes and tumour determined. Density plot show CD25 vs Foxp3-EGFP, and CD62L vs CD44 used to gate naive and EM phenotype Treg. Plots of CD45.1 and CD45.2 indicate host and donor composition of the indicated Treg subsets, and summary bar charts of normalised donor fraction across all mice (n=9 pooled from three independent experiments). (D) Foxp3FR mice were treated with a single feed of tamoxifen and 14 days later, engrafted with either A20 or EL4 tumour cells. (Foxp3-FR x Balb/c)F1 mice were used as hosts for A20 tumours. Mice were culled when tumour size reached maximal permitted size and phenotype of Treg assessed as in (B). Data are pool of 5 (B), 3 (C-D) independent experiments.

Finally, we asked whether this ontogeny of Treg might be model-specific, or if it rather represented a more generalised feature of Treg biology in malignant disease. We therefore analysed Treg infiltrates in two other model tumour systems, EL4 (thymoma) and A20 (B cell malignancy). Foxp3FR hosts were treated with TAM and then engrafted with tumours. After 2-3 weeks, tumours were recovered and their infiltrates analysed. In both cases, Treg recovered from the TME were EM phenotype, with elevated Ki67, and expressed the mTom reporter at a level that correlated closely with circulating EM Treg in the same host. These observations demonstrate again that the tumour-infiltrating Treg derived from cells that had already committed to a regulatory phenotype.

## Discussion

In this study, we wanted to define the ontogeny and phenotypic heterogeneity of circulating Treg, and to identify the pathways of Treg recruitment in malignant disease. Using independent fate reporter systems to track development and differentiation of Treg, our results strongly support a linear model of Treg development driven by self recognition. Thymus-derived Treg continually replenish circulating naive Treg throughout life, and these in turn differentiate into EM Treg constitutively throughout life, driven by self recognition. We also found strong evidence that tumour-infiltrating Treg derive specifically from circulating EM Treg.

Given the deleterious impact of immune suppression of beneficial anti-tumoural responses (Peng et al., 2012; Saleh and Elkord, 2020; Schnellhardt et al., 2022), the origin of regulatory T cells recruited to tumours has been a subject of intense study in recent years. Studies that suggest a significant contribution of inducible Treg to intratumour infiltrates have largely relied on correlative data, using markers or transcriptional profiles thought to distinguish iTreg and tTreg (Thornton et al., 2010; Weiss et al., 2012), or the similarity or difference of TCR repertoires between recirculating Treg and Treg from the TME (Savage et al., 2014). These approaches can be problematic since markers such as Nrp1 do not always correlate well with the apparent origin of Treg (Szurek et al., 2015). Further, the TCR repertoires of populations within the TME will inevitably be shaped by the activation events driving their recruitment and retention, making comparisons with putative precursor populations difficult. Our study employed genetic labelling of Foxp3^+^ Treg prior to engraftment in three independent tumour models, with the clear finding that reporter expression by Treg in the TME matched that of circulating Treg. While we cannot completely exclude a contribution of de novo generation of iTreg in these models, any contribution would at best represent a small fraction of the Treg in TME.

Our data also indicate that self recognition is the defining event controlling peripheral Treg homeostasis after thymic selection. The role of agonist selection in generating thymic Treg is well recognised. However, the antigens driving development of circulating antigen-experienced peripheral Treg are less well characterised. Studies using model antigens and TCR transgenic T cells have highlighted the importance of mucosal antigens, including microbiota, in driving development of RORgt-expressing Treg (Akagbosu et al., 2022; Kedmi et al., 2022; Weiss et al., 2012). However, these cells preferentially migrate to the GI tract, are only present as a small fraction of Treg in mesenteric lymph nodes that drain the intestines, and are not present in other lymph nodes or the spleen. Our analysis of WT mice and busulfan chimeras show that EM-Treg are generated continuously throughout life. Although the circulating Treg compartment is dominated by thymus-derived naive Treg early in life, EM Treg numbers increase steadily with age, coincident with a decline in thymic output. In busulfan chimeras, we saw almost half the EM Treg compartment is replaced by newly generated EM Treg by ∼d160 post transplant, and the kinetics of *Cre* reporter expression in Foxp3FR mice indicates these cells derive from reporter-labeled naive Treg precursors. Strikingly, this tonic generation appears to occur independently of microbiota, since the kinetics of de novo production of EM Treg were identical in “dirty” facilities that were sufficient to induce substantial inflation of conventional memory T cell subsets (Hogan et al., 2019). Given that thymically derived naive Treg are selected on self antigens, it seems reasonable to conclude that EM Treg development is also driven by a peripheral self-recognition event. However, it appears that only a subset of the naive Treg repertoire has the capacity to develop into EM Treg. This heterogeneity may reflect differences in either the diversity or abundance of self antigens expressed in peripheral lymphoid tissues, as compared with the thymus, where self antigen expression by medullary thymic epithelial cells is facilitated by AIRE (Peterson et al., 2008). There have, however, been reports of AIRE-expressing ILC3 in lymph nodes (Yamano et al., 2019), that may represent an additional source of self antigens that promote EM Treg development from naive Treg precursors in peripheral compartments. The specificity of EM Treg is especially pertinent when considering intratumoral Treg, since our results strongly indicated that Treg in the TME derive specifically from pre-existing EM Treg. This suggests that Treg, already primed by self antigens in the periphery, are preferentially recruited to tumours. This is an important distinction from the possibility that neoantigens drive activation and recruitment from either naive Treg or inducible Treg from conventional T cells.

Our computational analysis of Treg cell dynamics provided novel insights into the cellular mechanisms responsible for maintaining circulating naive and EM Treg. Naive Treg, like their conventional counterparts (Hogan et al., 2015), demonstrated a reliance on thymic output throughout life. Their numbers decline slowly with age, mirroring the atrophy of the thymus. However, in contrast to conventional naive T cells, we found evidence that self-renewal also made a substantial contribution to maintaining naive Treg numbers, exceeding that of thymic output as mice age; as a result, Treg represent 10-15% of the total CD4^+^ T cell compartment, even though only ∼1% of CD4SP thymocytes give rise to Treg. The additional contribution of self-renewal means that naive Treg TCR clonotypes may be larger on average than their conventional naive counterparts.

Previous studies have suggested that Treg recirculating to the thymus exert negative feedback upon thymic Treg development (Thiault et al., 2015). Our estimates of newly produced Treg as a fraction of total thymic Foxp3^+^ T cells were in close agreement with these earlier results, validating these observations. However, there were two confounding issues that contrived to exaggerate the apparent decline in thymic Treg development. First was the accumulation of EM Treg in thymi, which reached a maximal level already by 60 days of age and remained stable thereafter. Second was the coincident natural thymic atrophy that progressively reduces the rate of de novo generation of thymic Treg, because of dwindling precursors. Relating de novo Treg numbers instead to the size of the upstream precursor pool, CD4SPs, revealed stable levels of de novo Treg development for the first 8 months, with evidence of modest reduction by one year of age. The kinetics of recirculated EM Treg accumulation and reduction in Treg development do not correlate well. Given the age-related changes that take place during thymic atrophy (Liang et al., 2022), it is possible that the modest reduction in Treg output may instead reflect alterations in the function of some other component of thymic function such as mTECs, that are important for Treg development (Cowan et al., 2013). Analysis of donor infusion of peripheral naive Treg pools in busulfan chimeras did, however, exhibit a clear ceiling on the extent of replenishment possible, at around 0.75. A reduced thymic output of naive Treg with age could account for this observation, but would require an almost complete cessation of output by around 100-150d post BMT, a point at which we could still detect de novo Treg in abundance. In line with this, a simple homogeneous model of naive Treg homeostasis, which took the waning levels of de novo Treg in the thymus as input, still predicted replenishment approaching 90%. Instead, we suggest that the naive Treg pool is homeostatically heterogeneous at the time of bone marrow transplant, comprising a subset of Treg with enhanced survival and self renewal properties that confer a competitive advantage over newly generated thymic Treg. The mechanisms resulting in the generation and maintenance of such a subset remain to be determined, but given the rapid expansion of both Treg and conventional pools that occurs in neonatal mice, it is possible that Treg that develop during this window exhibit enhanced homeostatic properties to facilitate the rapid population of the peripheral pool.

Finally, the present study and our previous work provide clues regarding the functional importance of peripheral Treg activation states and dynamics. First, our conclusion here that EM Treg are constitutively generated throughout life by self recognition events, parallels our earlier studies of conventional CD4 memory T cell dynamics. In those studies we showed that central and CD4^+^ EM T cells are continuously generated and retained in healthy unchallenged laboratory mice, independently of microbiota and foreign antigenic stimulation (Gossel et al., 2017; Hogan et al., 2019)(Bullock et al 2024, in press). It is striking that continuous generation of putatively auto-reactive conventional CD4^+^ EM T cells is not also associated with development of overt autoimmune pathology. We speculate that the tonic generation of EM Treg that occurs in parallel alongside conventional CD4^+^ memory subsets represents an active and necessary regulatory response. It also seems plausible that the two processes share antigenic drivers. IL-2 production by conventional T cells is essential for normal Treg survival and homeostasis (Owen et al., 2018) so it is possible that Treg rely upon IL-2 produced during the constitutive generation of conventional CD4^+^ effector T cells. Second, we reveal the pathological consequences of maintaining a pool of recirculating self reactive EM Treg. Our work strongly indicates that Treg that first infiltrate the TME derive predominantly from circulating EM Treg, and not from conventional T cells or from the activation of naive Treg. While there is much interest in the role of tumour neo-antigens in shaping both conventional and regulatory T cell responses to tumours, the recruitment of self-reactive EM Treg to the TME would appear the most teleological solution for tumours to generate the most suppressive environment, since self antigens likely remain the most abundant source of regulatory T cell stimulation, regardless of tumour evolution and adaptation. An appreciation of the ontogeny of Treg found in TME may be essential if they are to be successfully targeted by immunotherapy without compromising self tolerance.

## Materials and methods

### Busulfan Chimeras and Foxp3 reporter strains

Busulfan chimeric C57Bl6/J mice were generated as described in (ref Hogan bio protocols). In summary, B6.SJL-*Ptprc^a^ Pepc^b^*/BoyJ mice (B6.CD45.1) and C57Bl6/J (B6.CD45.2) mice were bred and maintained in conventional colonies in the Comparative Biology Unit, Royal Free Campus of University College London, or where indicated, at the MRC National Institute for Medical Research, London, UK (NIMR). At NIMR, mice were housed in open cages and fed tap water. At UCL, mice were housed in individually ventilated cages, fed irradiated food and drank irradiated water. B6.CD45.1 mice aged between 8 and 25 weeks were treated with 20 mg/kg busulfan (Busilvex, Pierre Fabre) to deplete HSC, and reconstituted with T-cell depleted bone marrow cells from congenic donor B6.CD45.2 mice. Chimeras were sacrificed at various times after bone marrow transplantation. Cervical, brachial, axillary, inguinal and mesenteric lymph nodes, spleen and thymus were dissected from mice; single cell suspensions prepared, and analysed by flow cytometry.

*Foxp3^tm9(EGFP/cre/ERT2)Ayr^*/J mice (Jax strain 016961, *Foxp3^EGFP-CreERT2^*)(ref) and B6.Cg-*Gt(ROSA)26Sor^tm9(CAG-tdTomato)Hze^*/J mice (Jax strain 7909, *Rosa26R^mTom^* hereon) were obtained from Jax Laboratories and interbred to homozygosity at both loci. Mice were fed with tamoxifen by a single feed of 2mg of tamxoxifen (Sigma) diluted in 100µl corn oil (Fisher Scientific).

All experiments were performed in accordance with UK Home Office regulations, project license number PP2330953.

### Cell lines and tumour engraftment

MC38 (colon carcinoma), EL4 (T cell lymphoma) and A20 (B cell lymphoma) cell lines were passaged in vitro using standard conditions in DMEM culture media with 10% FCS. MC38 cells were passaged using 0.5% Trypsin EDTA 1X solution (Gibco). For engraftment into mice, cells were recovered at the exponential phase of growth in vitro, washed and resuspended in tissue culture grade 1X Phosphate Buffer Saline (PBS). Cells were then counted using the CASY counter and injected (5 × 10^6^/host) via the s.c. route, near the neck fold of host mice. Body weight and tumour diameter measurements were done at regular intervals. Tumours were permitted to grow up to a maximal diameter of 15mm over a period of 14-20 days. A20 tumours were transplanted into male (male Balb/c x female *Foxp3^EGFP-CreERT2^ Rosa26R^mTom^*)F1 mice.

### Flow cytometry and electronic gating strategies

Flow cytometric analysis was performed with 2-5 × 10^6^ thymocytes, 1-5 × 10^6^ lymph node or spleen cells. Cell concentrations of thymocytes, lymph nodes and spleen cells were determined with a Scharf Instruments Casy Counter. Cells were incubated with saturating concentrations of antibodies in 100 μl of Dulbecco’s phosphate-buffered saline (PBS) containing bovine serum albumin (BSA, 0.1%) for 1hour at 4°C followed by two washes in PBS-BSA. Cells were stained with the following monoclonal antibodies and cell dyes: CD45.1 FITC, CD45.2 AlexaFluor 700, TCR-beta APC, CD4 PerCP-eFluor710, CD25 PE, CD44 APC-eFluor780, CD25 eFluor450, CD62L eFluor450 (all eBioscience), TCR-beta PerCP-Cy5.5, CD5 BV510, CD4 BV650, CD44 BV785 (all BioLegend), CD62L BUV737 (BD Biosciences), LIVE/DEAD nearIR and LIVE/DEAD Blue viability dyes (Invitrogen). Foxp3 and Ki67 co-staining was performed using the FITC Flow Kit (BD Biosciences) according to the manufacturer’s instructions, along with anti-Ki67 eFluor660 (eBioscience). Cells were acquired on a BD LSR-II or BD 605 LSR-Fortessa flow cytometer and analysed using Flowjo software (Treestar). Subset gates were as follows: CD4 naive: live TCRβ^+^ CD4^+^ Foxp3^-^ CD44^lo^ CD62L^hi^. CD4 TEM: live TCRβ^+^ CD4^+^ Foxp3^-^ CD44^hi^ CD62L^lo^. CD4 TCM: live TCRβ^+^ CD4^+^ Foxp3^-^ CD44^hi^ CD62L^hi^. Naive Treg: live TCRβ^+^ CD4^+^ Foxp3^+^ CD44^lo^ CD62L^hi^. Effector memory Treg: live TCRβ^+^ CD4^+^ Foxp3^+^ CD44^hi^ CD62L^lo^.

### Mathematical modelling and statistical analysis

The mathematical models and our approach to model fitting are detailed in Text S1 of in SI. All code and data used to perform model fitting, and details of the prior distributions for parameters, are available at https://github.com/sanketrane/Treg_dynamics. Models were ranked using the Leave-One-Out (LOO) cross validation method (Vehtari et al., 2017). We quantified the relative support for models with the expected log point-wise predictive density (ELPD), for which we report the standard error. Models with ELPD differences below 4 were considered similar.

### Statistics

Statistical analysis, line fitting, regression analysis, and figure preparation were performed using Graphpad Prism 8. Column data compared by unpaired Mann-Witney student’s t test. * p<0.05, ** p<0.01, *** p<0.001, **** p < 0.0001, unless otherwise stated.

## Acknowledgements

We thank UCL Comparative Biology Unit staff for assistance with mouse breeding and maintenance. The authors declare no competing financial interests.

