## Supplementary material for "A linear ontogeny accounts for the development of naive, memory and tumour-infiltrating regulatory T cells in mice": SI Material

### S1 Mathematical modeling

#### S1A Models of naive Treg homeostasis

We modeled the dynamics of naive Tregs in busulfan chimeric mice – the changes in their pool size, their donor:host composition and the extent of their Ki67 expression – using a mechanistic modeling approach. First, we considered a simple **homogeneous model**, which assumes that the birth, death, and maturation dynamics of naive Tregs are governed by processes that remain unchanged over the mouse's lifetime. This model can only explain the incomplete replacement (*i.e.,* normalised chimerism stabilizing to values < 1) in naive Tregs that we observe when the influx of precursor-derived cells declines with time and drops below their net loss rate. That is, the supply of donor-derived cells from their precursors dries up before their donor fraction can equilibrate with the chimerism in their ancestors in the thymus – the early stage CD4 CD8 double positive (DP) thymocytes.

Alternatively, the partial replacement of host cells by donor-derived cells in naive Tregs can also be explained by models assuming heterogeneity in their maintenance. To address this, we included the **incumbent model** in our analyses, which compartmentalizes naive Treg population into two subsets with distinct kinetics birth and loss. Specifically, it defines an 'incumbent' subset composed of self-renewing, long-lived subpopulation of relatively older cells that are established early on in life – before the bone marrow transplant – which are resistant to replacement by newly derived donor cells. The incumbent model also considers a 'displaceable' subset within naive Tregs consisting cells with comparatively rapid kinetics of division and loss. For the purposes of modeling naive Treg dynamics in adult (7 weeks and older) busulfan chimeric mice, we consider that thymic precursors of naive Tregs exclusively feed into the displaceable subset.

##### General formulations of the neutral and incumbent models

We defined both of these models as the systems of ordinary differential equations (ODE), to separately track the dynamics of incumbent and displaceable compartments in Ki67$^+$ and Ki67$^-$ subsets of donor- and host-derived naive Treg cells.

**Neutral model:**

Host compartment in busulfan chimeras

Ki67$^+$subset

$$\frac{dX_{\mathsf{h}}^{+}}{dt} = \alpha\,\phi(t)\,(1-\chi(t))\,\epsilon_h + \rho_{\mathsf{x}}\,(2\,X_{\mathsf{h}}^{-} + X_{\mathsf{h}}^{+}) - (\beta+\delta_{\mathsf{x}})\,X_{\mathsf{h}}^{+}$$

Ki67$^-$subset

$$\frac{dX_{\mathsf{h}}^{-}}{dt} = \alpha\,\phi(t)\,(1-\chi(t))\,(1-\epsilon_h) + \beta\,X_{\mathsf{h}}^{+} - (\rho_{\mathsf{x}}+\delta_{\mathsf{x}})\,X_{\mathsf{h}}^{+}$$

(1)

Donor compartment in busulfan chimeras

Ki67$^+$subset

$$\frac{dX_{\mathsf{d}}^{+}}{dt} = \alpha\,\phi(t)\,\chi(t)\,\epsilon_d + \rho_{\mathsf{x}}\,(2\,X_{\mathsf{d}}^{-} + X_{\mathsf{d}}^{+}) - (\beta+\delta_{\mathsf{x}})\,X_{\mathsf{d}}^{+}$$

Ki67$^-$subset

$$\frac{dX_{\mathsf{d}}^{-}}{dt} = \alpha\,\phi(t)\,\chi(t)\,(1-\epsilon_d) + \beta\,X_{\mathsf{d}}^{+} - (\rho_{\mathsf{x}}+\delta_{\mathsf{x}})\,X_{\mathsf{d}}^{+}$$

We used empirical descriptions of the total size of the immediate precursors of naive Tregs and the extent of donor chimerism and fraction of Ki67$^+$ cells within the precursors to define $\phi(t), \chi(t)$ and $\epsilon(t)$. These descriptor functions are explained in detail in the **section S1C**. Both the neutral (equation 1) and the incumbent (equation 2) model estimate the rate of daily influx ($\alpha$) into naive Tregs, their division and loss rates ($\rho$ and $\delta$) using the data from the busulfan chimeras. Lastly, we used our previous estimate of the rate of loss of Ki67 expression ($\beta$) on T cells (Gossel et al eLIfe) and fixed it to $1/3.5$ days$^{-1}$. We depict the general formulation of the incumbent model similarly below.

**Incumbent model:**

Host compartment in busulfan chimeras

Displaceable Ki67$^+$subset

$$\frac{dX_\text{h}^+}{dt} = \alpha\,\phi(t)\,(1-\chi(t))\,\epsilon(t) + \rho_\text{x}\,(2\,X_\text{h}^- + X_\text{h}^+) - (\beta + \delta_\text{x})\,X_\text{h}^+$$

Displaceable Ki67$^-$subset

$$\frac{dX_\text{h}^-}{dt} = \alpha\,\phi(t)\,(1-\chi(t))\,(1-\epsilon(t)) + \beta\,X_\text{h}^+ - (\rho_\text{x} + \delta_\text{x})\,X_\text{h}^+$$

Incumbent Ki67$^+$subset

$$\frac{dY_\text{h}^+}{dt} = \rho_\text{y}\,(2\,Y_\text{h}^- + Y_\text{h}^+) - (\beta + \delta_\text{y})\,Y_\text{h}^+$$

Incumbent Ki67$^-$subset

$$\frac{dY_\text{h}^-}{dt} = \beta\,Y_\text{h}^+ - (\rho_\text{y} + \delta_\text{y})\,Y_\text{h}^+ \tag{2}$$

Donor compartment in busulfan chimeras

Displaceable Ki67$^+$subset

$$\frac{dX_\text{d}^+}{dt} = \alpha\,\phi(t)\,\chi(t)\,\epsilon(t) + \rho_\text{x}\,(2\,X_\text{d}^- + X_\text{d}^+) - (\beta + \delta_\text{x})\,X_\text{d}^+$$

Displaceable Ki67$^-$subset

$$\frac{dX_\text{d}^-}{dt} = \alpha\,\phi(t)\,\chi(t)\,(1-\epsilon(t)) + \beta\,X_\text{d}^+ - (\rho_\text{x} + \delta_\text{x})\,X_\text{d}^+$$

Incumbent Ki67$^+$subset

$$\frac{dY_\text{d}^+}{dt} = \rho_\text{y}\,(2\,Y_\text{d}^- + Y_\text{d}^+) - (\beta + \delta_\text{y})\,Y_\text{d}^+$$

Incumbent Ki67$^-$subset

$$\frac{dY_\text{d}^-}{dt} = \beta\,Y_\text{d}^+ - (\rho_\text{y} + \delta_\text{y})\,Y_\text{d}^+$$

#### S1B Modeling EM Treg dynamics

We used a two-compartmental model that encodes heterogeneity in EM Treg turnover to explain their behaviour in busulfan chimeric mice. We argued that activation and cell-division events accompany differentiation to the EM Treg lineage, and therefore designed a model that divides cells into two kinetically distinct subsets based on the time since their compartmental entry. In this **two-compartment** model, we considered that new precursor-derived cells enter EM Treg compartment via a 'fast' subset with relatively rapid dynamics of division and death. Cells in the fast before transitioning into a more quiescent 'slow' state.

**ODE system depicting the two-compartment model:**

Host compartment in busulfan chimeras

Fast Ki67$^+$subset

$$\frac{dX_h^+}{dt} = \alpha\,\phi(t)\,(1-\chi(t)) + \rho_x\,(2\,X_h^- + X_h^+) - (\beta + \delta_x + \mu)\,X_h^+$$

Fast Ki67$^-$subset

$$\frac{dX_h^-}{dt} = \beta\,X_h^+ - (\rho_x + \delta_x + \mu)\,X_h^+$$

Slow Ki67$^+$subset

$$\frac{dY_h^+}{dt} = \mu\,X_h^+ + \rho_y\,(2\,Y_h^- + Y_h^+) - (\beta + \delta_y)\,Y_h^+$$

Slow Ki67$^-$subset

$$\frac{dY_h^-}{dt} = \mu X_h^+ + \beta\,Y_h^+ - (\rho_y + \delta_y)\,Y_h^+$$

(3)

Donor compartment in busulfan chimeras

Fast Ki67$^+$subset

$$\frac{dX_d^+}{dt} = \alpha\,\phi(t)\,(1-\chi(t)) + \rho_x\,(2\,X_d^- + X_d^+) - (\beta + \delta_x + \mu)\,X_d^+$$

Fast Ki67$^-$subset

$$\frac{dX_d^-}{dt} = \beta\,X_d^+ - (\rho_x + \delta_x + \mu)\,X_d^+$$

Slow Ki67$^+$subset

$$\frac{dY_d^+}{dt} = \mu\,X_d^+ + \rho_y\,(2\,Y_d^- + Y_d^+) - (\beta + \delta_y)\,Y_d^+$$

Slow Ki67$^-$subset

$$\frac{dY_d^-}{dt} = \mu X_d^+ + \beta\,Y_d^+ - (\rho_y + \delta_y)\,Y_d^+$$

Here, we considered that all precursor-derived cells enter the Ki67$^+$ subset, since cell-division is a requisite for the commitment to the EM Treg lineage. We estimated the rates of influx ($\alpha$), maturation of fast to slow subest ($\mu$), division ($\rho$) and loss ($\delta$), using the data-derived from busulfan chimeras. Additionally, we fixed the rate of loss of Ki67 expression $\beta = 1/3.5$ days$^{-1}$, similar to as described for the neutral and incumbent models (eq. 1 and 2).

#### S1C Modeling the de novo production, division and loss of naive and EM Treg

Our modeling strategy allows us to map the chimerism and Ki67 expression kinetics of the potential source population onto the dynamics of the population of interest. Specifically, the chimerism in the source constrains the parameters defining influx and 'net loss' (death - division), while the Ki67$^+$ fraction of the precursor population constrains influx and division rates of the population of interest. By comparing the quality of the model fits we then established the relative support for different candidate precursors of Tregs.

##### Precursors of naive and EM Tregs

We considered that *de novo* generated FoxP3$^+$ thymic SP4 T cells are direct precursors of naive Tregs. For EM Tregs, we compared naive Tregs and conventional naive, central memory, and effector memory CD4 T cells as potential precursors. We describe the changes in their total pool size an donor:host chimerism using phenomenological functions described below. We use the age of the youngest animal in busulfan chimera set as the starting point for modeling *i.e.,* $t_0 = 40$ days.

##### Describing influx into the Treg compertments

We used a general descriptor function, described in **equation 4**, to capture the changes in the pool sizes of precursor populations of naive and EM Tregs.

$$\phi(t) = \phi_0 \, e^{-\frac{(t-t_0)}{p}}. \tag{4}$$

Our model definitions assume that rate of influx into both naive and EM Tregs '$\alpha$' remains unvarying with time and is proportional to the size of the precursor population under consideration. The rate of precursor influx is then $\alpha \times \phi(t)$, where the constant $\alpha$ is estimated while fitting models to the busulfan chimera data. We estimated the parameters $\phi_0$ and $\nu$ by fitting **equation 4** to the log-transformed counts of the precursor population. We show the descriptions of pool size changes and parameter estimates for the precursors of naive Tregs *i.e., de novo* generated FoxP3$^+$ thymic SP4 T cells in **Fig. S1A**. We show the descriptions of pool sizes for the candidate precursors of EM Tregs *viz.* the conventional naive, central memory, and effector memory CD4 T cells in **Fig. S2A**.

##### Empirical descriptions of normalised chimerism in the precursor populations

We describe the changes in the fraction of donor cells in precursors of naive and EM Tregs using

$$\chi(t) = \chi_0 \left(1 - e^{-\frac{(t-t_0)^n}{q}}\right) \tag{5}$$

The parameters $\chi_0$ and $q$ were estimated by fitting **equation 4** to the observed time-course of donor chimerism in the precursor population in busulfan chimeras. Fits and parameter estimates for the precursor populations are shown in **Fig. S1B** and **S2B**.

##### Ki67 expression dynamics in the precursors of naive and EM Tregs

The fractions of Ki67 expressing cells in both the host- and donor-derived FoxP3$^+$ thymic SP4 T cells (mean values 15% and 9%, respectively) remain stable across mouse lifetime **Fig. S1C**. We assumed that these dynamics are represented in the influx into naive Tregs and fixed the rate constants $\epsilon_h$ and $\epsilon_d$ that defined Ki67$^+$ fraction in precursor influx to 0.15 and 0.09, in **equations 1** and **2.**

We assumed that all newly generated EM Tregs were Ki67$^+$ , since differentiation into memory likely involves cell division.

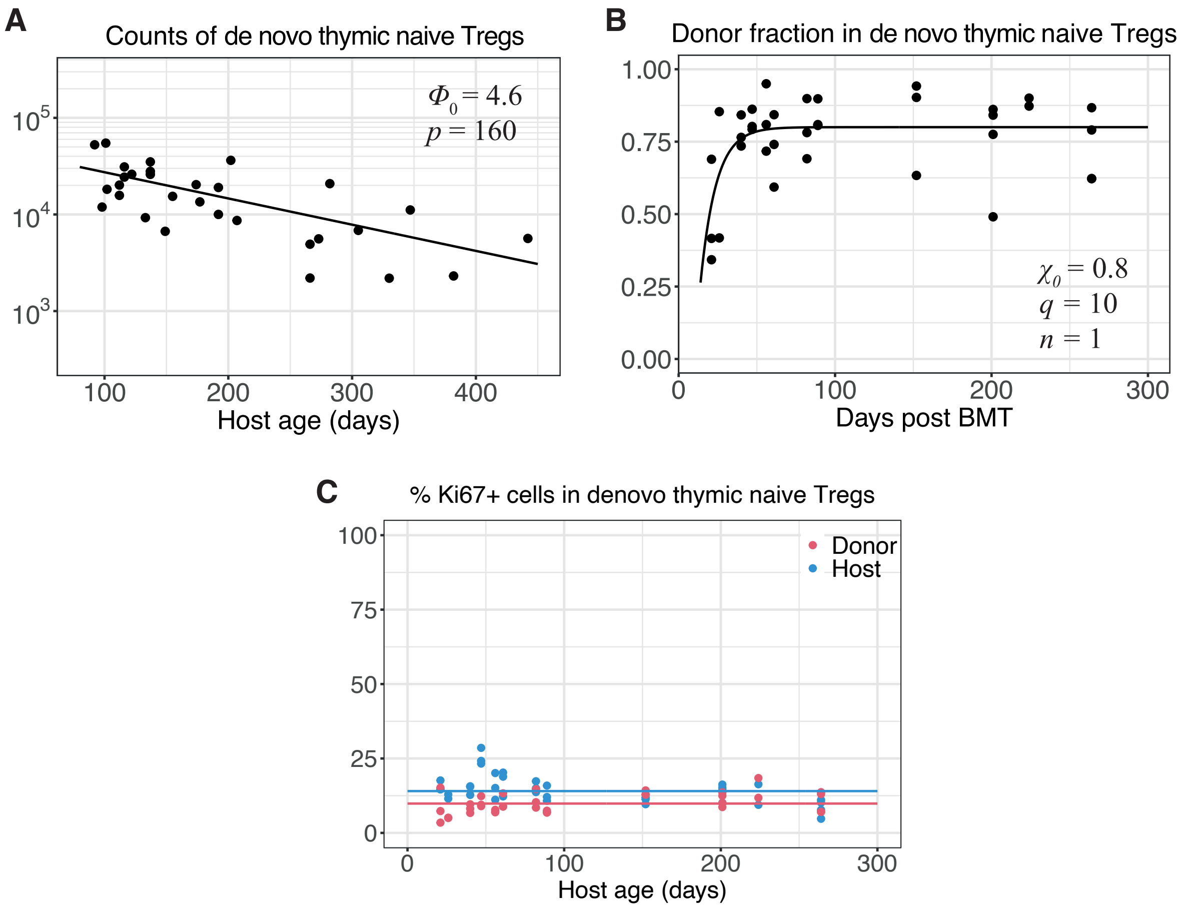


**Figure S1: Dynamics of naive Treg precursors.** We show the changes in pool size **(A)**, donor chimerism **(B)** and Ki67$^+$ fraction **(C)** of *de novo* generated FoxP3$^+$ thymic SP4 T cells. Curves shown in **(A)** and **(B)** were generated using **equation 4** and **equation 5**, respectively. The intercepts shown in **(C)** were fixed to the mean Ki67$^+$ percentages observed among host and donor FoxP3$^+$ thymic SP4 T cells – 15% and 9%, respectively.

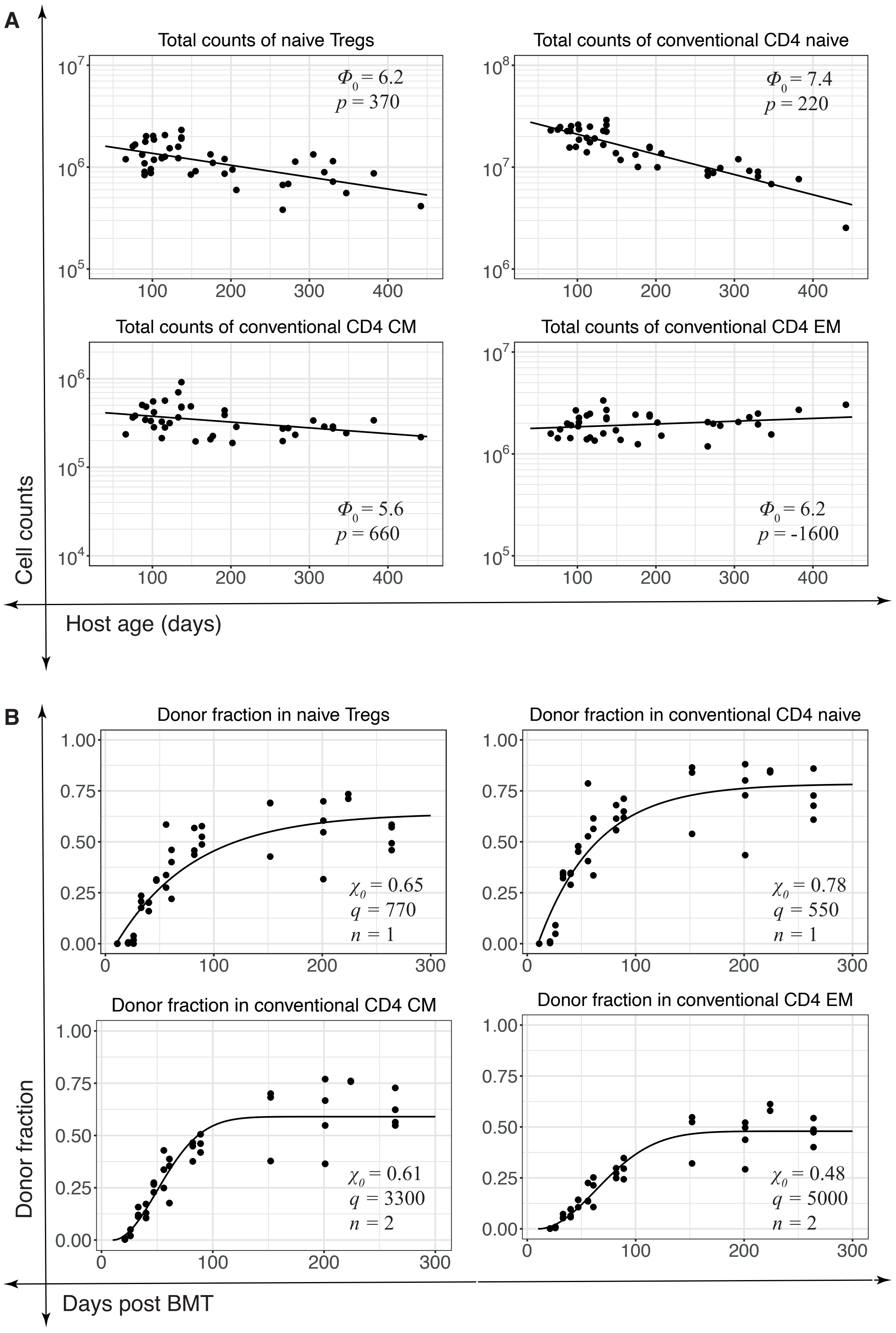


**Figure S2: Dynamics of potential precursor populations of EM Tregs.** We depict the time-courses of total counts **(A)** and donor chimerism **(B)** of candidate precursors of EM Treg cells. Lines shown **A** represent phenomenological descriptions of the observations of total counts (solid dots) generated using **equation 4**. Similarly, we generated curves for chimerism data in **B** using **equation 5**.

#### S1D Explaining the growth of EM Treg numbers over the first year of life

Our analysis suggests that EM Treg population contains a transient stage – the fast subset – which quickly transitions to a more persistent 'slow' stage with a mean time $\sim$ 5 days. The slow population divide and die on average every 40 days, but these processes are finely balanced - therefore, slow EM Treg accumulate over time as they are fed from the fast population. The growth in total EM Treg numbers during the first year of life can then be understood quite simply. If we sum Ki67$^+$ and Ki67$^-$ cells, and also sum host and donor cells, the following equations describe the timecourses of total numbers of fast ($X$) and slow ($Y$) EM Treg:

$$\frac{dX}{dt} = \alpha\,\phi(t) - (\lambda_X + \mu)X(t), \tag{6}$$

$$\frac{dY}{dt} = \mu\,X(t) - \lambda_Y Y(t), \tag{7}$$

where $\lambda_X = \delta_x - \rho_x$ and $\lambda_Y = \delta_y - \rho_y$ are the net rates of loss (that is, loss minus self-renewal) for fast and slow cells respectively. Adding these,

$$\frac{dM}{dt} = \alpha\,\phi_0 e^{-(t-t_0)/p} - \lambda_X X(t) - \lambda_Y Y(t), \tag{8}$$

where $M(t)$ is total EM Treg numbers, and we show explicitly the waning influx described empirically by the declining exponential (equation 4). Equation 8 shows that during the first year of life, when $X$ is small, the influx of new EM cells outstrips the total rate of loss fast cells. From equation 7 we see slow cells continuously accumulate during this period, since their net loss rate $\lambda_Y$ is close to zero. This accumulation explains the decline in Ki67 levels in bulk among EM Treg (Fig 4D), even while the fast cell population is increasing. Only after the influx of new EM declines sufficiently for fast cells to decline, would EM cell numbers start to fall; the model predicts this would happen in very old age, beyond the time frame of these experiments.

#### S2 Model validation and comparison

Each model in our modeling framework is fitted simultaneously to four sets of observations – cell counts ($y_{a,1}$), normalized donor fraction ($y_{a,2}$) and the fraction of Ki67$^+$ cells within donor ($y_{a,3}$) and host ($y_{a,4}$) cells, where $a$ denotes the animal (a = 1,…,n).

We estimate the model parameters using a Bayesian statistical inference approach. The joint density of the observations in each animal is defined as, $y_a = (y_{a,1}, y_{a,2}, y_{a,3}, y_{a,4})$. We assume that $y_a$ have independent multivariate normal distributions with mean $\mu_a = (\mu_{a,1}, \mu_{a,2}, \mu_{a,3}, \mu_{a,4})$ and covariance matrix $D = \text{diag}(\sigma_1^2, \sigma_2^2, \sigma_3^2, \sigma_4^2)$. Here, $\mu_{a,i} = f_i(time_a, \theta)$, is the model prediction for $i^{\text{th}}$ observation in $a^{\text{th}}$ animal. The 'prior' distributions of model parameters are defined based on existing knowledge regarding their values. Observations from the posterior distribution of $(\theta, D)$ are generated using the no-U-turn-sampler sampler in the *Stan* language, where parameters are sampled from the joint prior density following the Hamiltonian Monte Carlo algorithm.

***Model selection criteria:*** We estimate the expected log point-wise predictive density (elpd$^j$) for each model ($M_j$), which is the measure of its performance and out-of-sample prediction accuracy[1,2], using the leave-one-out (LOO) cross validation method. The LOO process estimates the probability density $P(y_i|y_{-i}, M_j)$ of the prediction of $i^{\text{th}}$ observation using the model $M_j$ fitted on the data with observation $i$ excluded. The elpd estimate is then the sum of the predictive densities of the LOO estimates of all $n$ observations in the data,

$$\widehat{\text{elpd}^j_{\text{loo}}} = \sum_{i=1}^{n} \text{elpd}^j_{\text{loo, i}} = \sum_{i=1}^{n} \log(P(y_i|y_{-i}, M_j)), \qquad \text{se}(\widehat{\text{elpd}^j_{\text{loo}}}) = \sqrt{\sum_{i=1}^{n} \left(\text{elpd}^j_{\text{loo, i}} - \text{elpd}^j_{\text{loo}}/n\right)^2}. \tag{9}$$

We use the estimates of elpd and its standard error (eq. 9) to calculate the relative support for each model as the model weight ($W$) and to rank them using the Pseudo-Bayesian model averaging method, implemented in the *loo* package in *R*.

$$W_j = \frac{\exp\left(\widehat{\text{elpd}^j_{\text{loo}}} - \frac{1}{2}\text{se}(\widehat{\text{elpd}^j_{\text{loo}}})\right)}{\sum_{j=1}^{J} \exp\left(\widehat{\text{elpd}^j_{\text{loo}}} - \frac{1}{2}\text{se}(\widehat{\text{elpd}^j_{\text{loo}}})\right)}. \tag{10}$$

As the elpd estimates are derived from LOO cross validation, $W$ is interpreted as the confidence in a model's ability to predict new data, relative to all the other models under consideration.

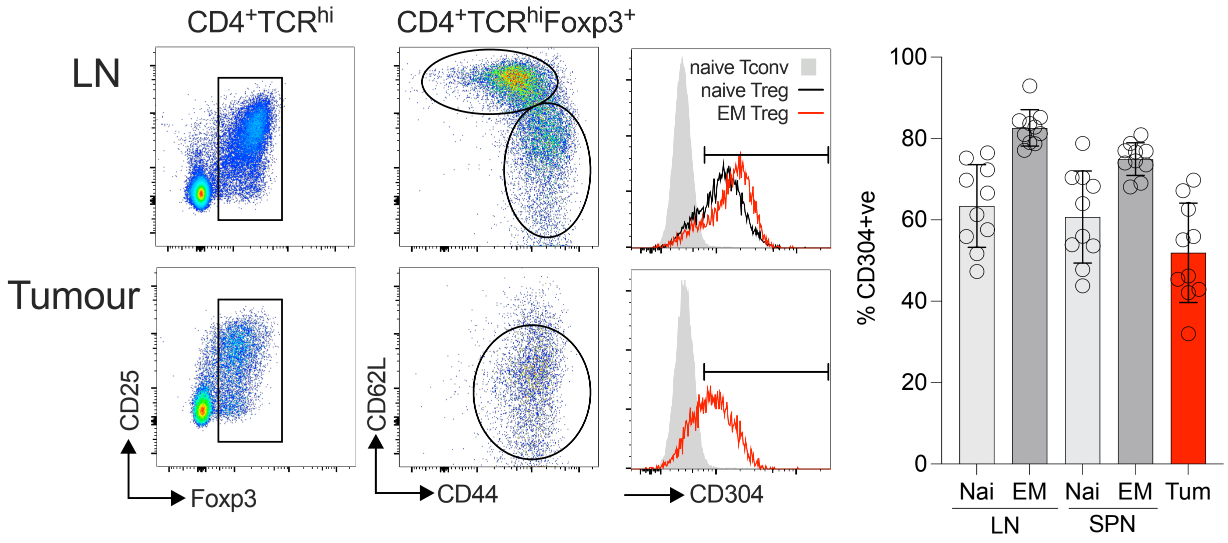


**Figure S3 - Nrp1 (CD304) expression by circulating and intratumoral Treg.** Treg from lymph nodes and tumours of busulfan chimeras described in figure 7C, engrafted with MC38 cells, were analysed for expression of CD304. Density plots show gating strategy to identify Foxp3+ Treg, and naive vs EM Treg subsets therein. Histograms show CD304 expression by naive and EM Treg from LN (as compared with naive CD4+ conventional T cells (TOP histogram), and EM Treg in tumour vs naive CD4+ conventional T cells from LN. Summary bar charts show % CD304+ in the indicated Treg subsets in LN, spleen and tumour (n=9 pooled from three experiments).
